# ME3BP-7 is a targeted cytotoxic agent that rapidly kills pancreatic cancer cells expressing high levels of monocarboxylate transporter MCT1

**DOI:** 10.1101/2023.07.23.550207

**Authors:** Jordina Rincon-Torroella, Marco Dal Molin, Brian Mog, Gyuri Han, Evangeline Watson, Nicolas Wyhs, Shun Ishiyama, Taha Ahmedna, Il Minn, Nilofer S. Azad, Chetan Bettegowda, Nickolas Papadopoulos, Kenneth W. Kinzler, Shibin Zhou, Bert Vogelstein, Kathleen Gabrielson, Surojit Sur

**Affiliations:** Ludwig Center, Sidney Kimmel Comprehensive Cancer Center, The Johns Hopkins University School of Medicine, Baltimore, MD 21287, USA; Department of Neurosurgery, The Johns Hopkins University School of Medicine, MD 21205, USA; Department of Oncology, The Johns Hopkins University School of Medicine, Baltimore, MD 21287, USA; Department of Surgery, The Johns Hopkins University School of Medicine, Baltimore, MD, USA; Howard Hughes Medical Institute, Chevy Chase, MD 20815, USA; Lustgarten Pancreatic Cancer Research Laboratory, Sidney Kimmel Comprehensive Cancer Center, The Johns Hopkins University School of Medicine, Baltimore, MD 21287, USA; Department of Biomedical Engineering, Johns Hopkins University, Baltimore, MD 21218, USA; Department of Molecular and Comparative Pathobiology, The Johns Hopkins University School of Medicine; Sidney Kimmel Comprehensive Cancer Center, Johns Hopkins University; Division of Nuclear Medicine and Molecular Imaging, The Russell H. Morgan Department of Radiology and Radiological Science, The Johns Hopkins University School of Medicine, 601 North Caroline Street, JHOC 3223, Baltimore, MD, 21287, USA; Bloomberg∼Kimmel Institute for Cancer Immunotherapy, Sidney Kimmel Comprehensive Cancer Center, Baltimore, MD 21287, USA; Department of Pathology, The Johns Hopkins University School of Medicine, Baltimore, MD 21205, USA

## Abstract

Nearly 30% of Pancreatic ductal adenocarcinoma (PDAC)s exhibit a marked overexpression of Monocarboxylate Transporter 1 (MCT1) offering a unique opportunity for therapy. However, biochemical inhibitors of MCT1 have proven unsuccessful in clinical trials. In this study we present an alternative approach using 3-Bromopyruvate (3BP) to target MCT1 overexpressing PDACs. 3BP is a cytotoxic agent that is known to be transported into cells via MCT1, but its clinical usefulness has been hampered by difficulties in delivering the drug systemically. We describe here a novel microencapsulated formulation of 3BP (ME3BP-7), that is effective against a variety of PDAC cells in vitro and remains stable in serum. Furthermore, systemically administered ME3BP-7 significantly reduces pancreatic cancer growth and metastatic spread in multiple orthotopic models of pancreatic cancer with manageable toxicity. ME3BP-7 is, therefore, a prototype of a promising new drug, in which the targeting moiety and the cytotoxic moiety are both contained within the same single small molecule.

**One Sentence Summary:** ME3BP-7 is a novel formulation of 3BP that resists serum degradation and rapidly kills pancreatic cancer cells expressing high levels of MCT1 with tolerable toxicity in mice.

## INTRODUCTION

Pancreatic ductal adenocarcinoma (PDAC) confers a dismal 5-year survival.^1,2^ Even after aggressive multimodality treatment, only a minority of patients achieve long term survival. The mainstay of treatment remains combinations of fluorouracil, irinotecan, oxaliplatin (FOLFIRINOX), or gemcitabine/nab-paclitaxel, which cause severe side effects and offer only modest improvements in survival^3–5^. Therefore, the development of new rationally-designed therapeutic agents are critical for improved outcomes in patients diagnosed with this devastating disease.

Metabolic reprogramming is one of the hallmarks of PDACs. Major effectors underlying the aberrant metabolic re-wiring in cancer are members of the monocarboxylate transporter (MCT) family. These transmembrane proteins mediate the transport of pyruvate, lactate, short-chain fatty acids, and ketones in and out of cells.^6–8^ Several studies have associated increased MCT1 expression in various cancers with chemoresistance and poor prognosis.^1,9–16^ Although MCT1 is an intriguing biological target, the small molecule MCT1 inhibitor AZD3965 was not found to be effective in the treatment of solid tumors. ^17,18^

3-Bromopyruvate (3BP) is a potent cytotoxic analog of pyruvic acid with a unique mode of action.^19^ The anti-cancer effects of 3BP were initially attributed to inhibition of glycolysis.^20–22^ However, recent studies have revealed 3BP as an alkylating agent of multiple intracellular proteins.^19,23^ MCT1 expression is essential for the activity of 3BP ^24^, although effective methods to systemically deliver 3BP have been a major research challenge. Clinical development of 3BP has been hampered by its poor serum stability, unfavorable pharmacokinetics, and excessive *in vivo* toxicity despite substantial pre-clinical and clinical efforts.^25,26^ The biochemical basis for these poor pharmacokinetic features is that 3BP alkylates free sulfhydryl groups in plasma proteins, including albumin, resulting in drug inactivation.^19^ To overcome this rapid loss of 3BP activity, large doses of the drug have been administered systemically or it was delivered locally to tumors, such as through the hepatic artery in case of liver tumors.^27^

Our previous work demonstrated that micro-encapsulation of 3BP reduces its toxicity^28^. Herein, we describe a novel encapsulated formulation of 3BP, ME3BP7, that reduces its degradation in serum. We show that ME3BP7 preferentially causes cell death in cells expressing MCT1 and inhibits the growth of orthotopic PDAC xenografts in mice without excessive toxicity.

## RESULTS

### MCT1 expression mediates sensitivity of PDAC cell lines to 3BP

Comparison of RNAseq datasets from TCGA and the Cancer Cell Line Encyclopedia (CCLE) with The Genotype-Tissue Expression (GTEx) portal revealed that ∼20-25% of PDACs exhibit increased expression of monocarboxylate transporter 1 (MCT1), a mediator of 3BP activity. We then explored the activity of 3BP as free drug in a panel of PDAC cell lines, including MIA PaCa-2, PSN-1, Panc 02.13, AsPC-1, BxPC-3, and CFPAC-1. Cells were treated with increasing concentrations of 3BP (0 to 220 µM) and assessed for cell death with real-time quantitative live-cell imaging. Five of the cell lines, MIA PaCa-2, PSN-1, Panc 02.13, AsPC-1, and BxPC-3, were sensitive to 3BP, with IC50s ranging from 24-40 µM (Fig. 1A and 1B). In contrast, CFPAC-1 was highly resistant to 3SP, at concentrations of up to 220 µM. RNA expression levels of three cellular transporters^29,30^, *GLUT1*, *MCT1* and *MCT4*, were examined to identify potential associations between the drug and cell viability (Fig. 1B). The expression of *MCT1* was much lower in the 3BP-resistant cell line CFPAC-1 than in the other five cell lines. However, no clear relationship was found between 3BP-resistance and the expression of the other transporters. MCT1 immunohistochemical (IHC) analyses furthermore demonstrated that the expression of the MCT1 protein on the cell surface paralleled RNA levels in these cell lines (Fig. 1C).

**Fig. 1.**
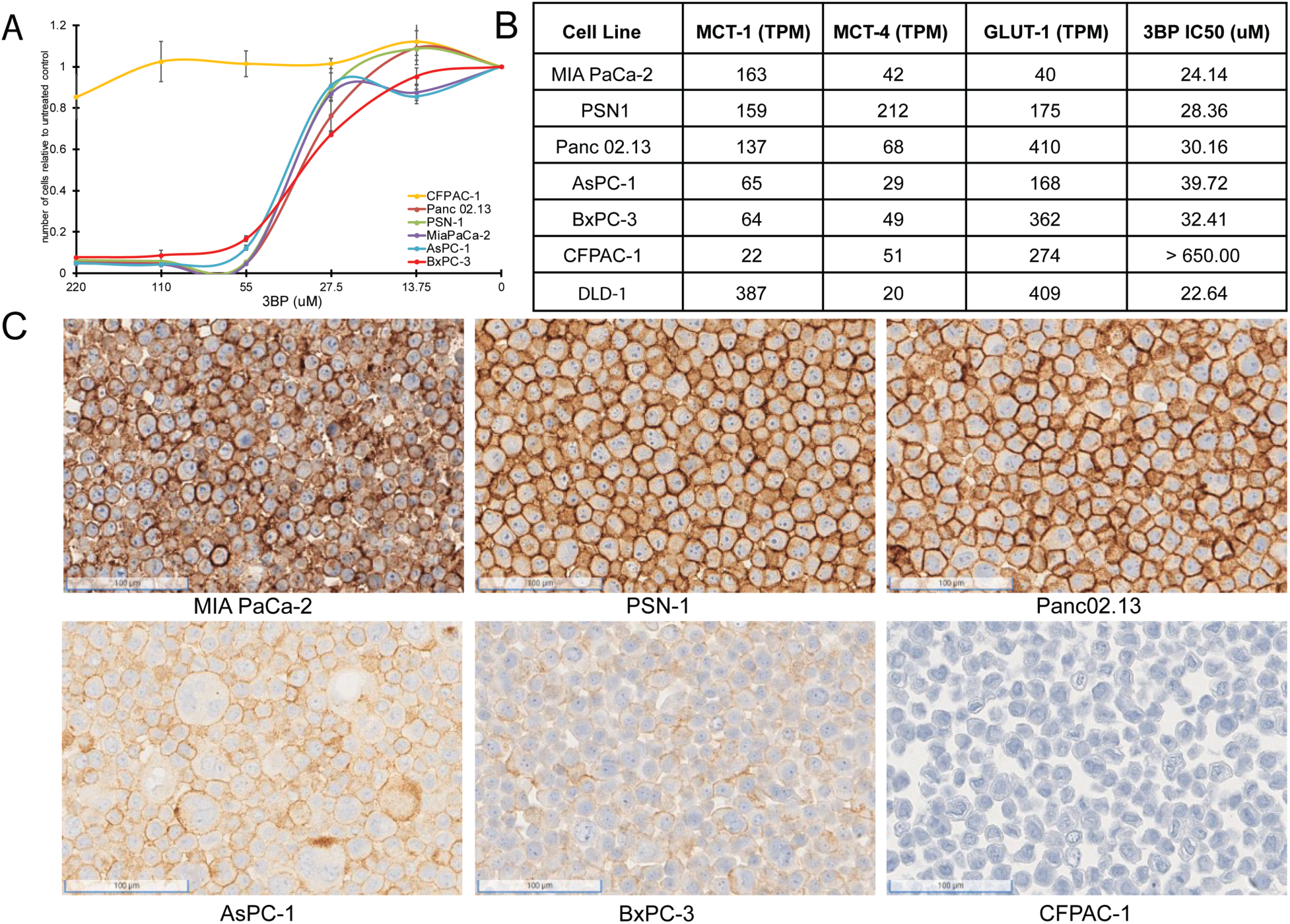
3BP sensitivity of pancreatic ductal adenocarcinoma cell lines. (**A**) Response of six different PDAC cell lines to 3-BP. The indicated cell lines were exposed to increasing doses of 3BP for 72 h and evaluated with a SYBR green growth assay. Data are represented as the mean +/- SD of three technical replicates and are normalized to untreated controls. (**B**) Expression levels of *MCT-1*, *MCT-4* and *GLUT-1* (TPM) and corresponding IC50s of 3BP. (**C**) Immunohistochemistry performed on PDAC cell lines with monoclonal mouse antibody against MCT1.

To further demonstrate specificity of the drug, we genetically inactivated MCT1 in cells and compared the viability of inactivated cells with their parental counterparts in the presence of 3BP. A similar strategy was used to establish MCT1 as essential for 3BP-mediated cell death in KBM7 cells, a unique near haploid cell line derived from a leukemia. ^24^ We deleted the gene encoding MCT1, *SLC16A1*, in the 3BP sensitive cell line MIA PaCa-2 expressing high levels of MCT1 (Fig. 1A and 1B). Next-generation sequencing confirmed knockout of MCT1 in six clones (Fig. 2A and 2B), and the results were validated with IHC (Fig. 2C and 2D). Cells from the six clones were then mixed at equal ratios to create an MCT1 KO cell population (KO cells), thereby avoiding potential variables often associated with the evaluation of a single clone. ^31,32^

**Fig. 2.**
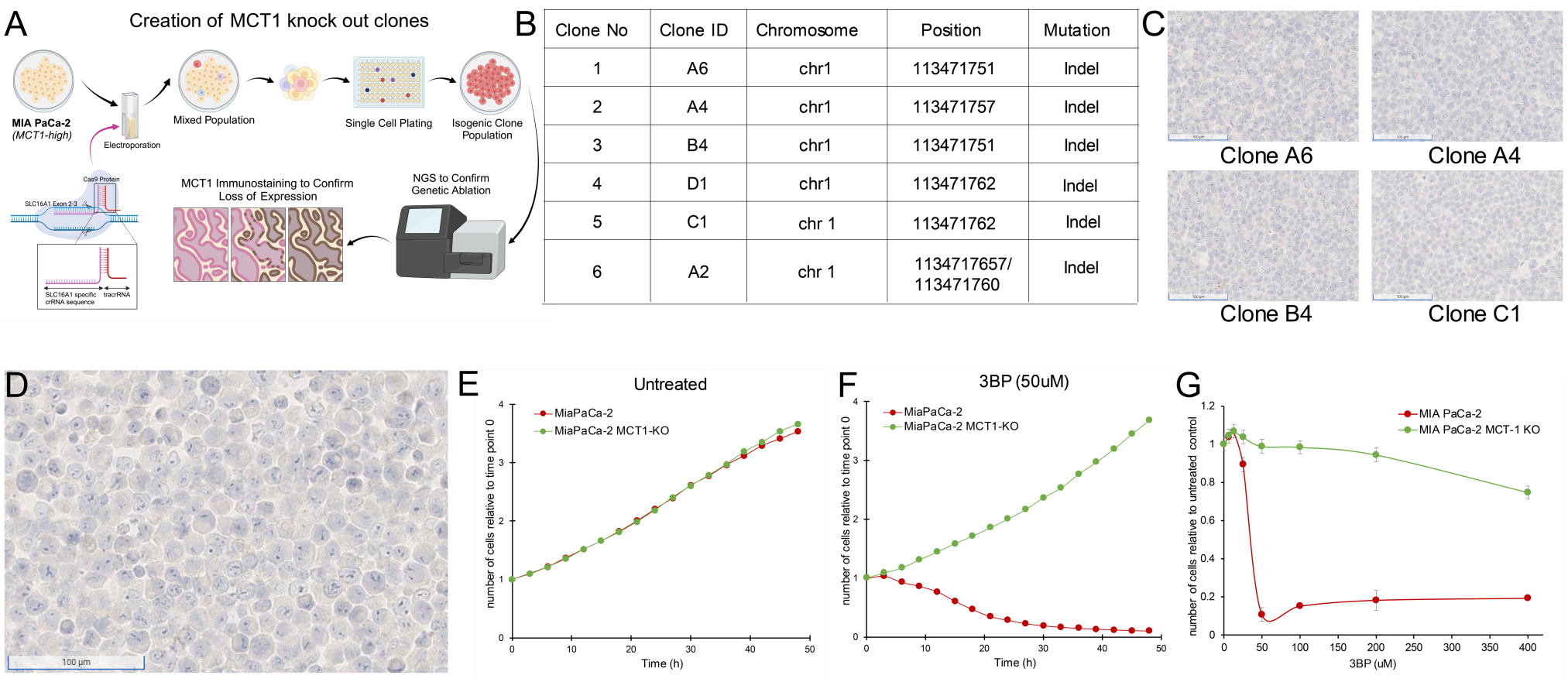
MCT-1 mediates 3BP activity. (**A**) Strategy for CRISPR based knockout of *SLC16A1* in MiaPaCa-2 cells. (**B**) Table of Knockout clones. (**C**) IHC performed on representative KO clones with monoclonal mouse antibody against MCT1 (1:2000 dilution). (**D**) IHC performed with MCT1 antibody on pooled monoclonal MCT1 KO cells used to test the MCT1 specific activity of 3BP and ME3BP-7. (**E** and **F**) Comparison of cell growth over time of MiaPaCa-2 and MiaPaCa-2 MCT1-KO in the absence and presence of 3BP (50 µM) normalized to time point 0 h. Data are represented as the mean +/- SD of two technical replicates. (**G**) Dose-response curves of MiaPaCa-2 cells and MiaPaCa-2 MCT1-KO at 36 h. Cell viability normalized to the number of cells at 0 h. Data represent the mean +/- SD of two technical replicates.

Next, we introduced nuclear-restricted green and red fluorescent proteins into the KO and parental cells, respectively, to enable simultaneous visualization in co-culture. KO cells grew at a rate indistinguishable from the parental cells under normal culture conditions (Fig. 2E). However, in the presence of 50 µM 3BP, the parental cells, with an IC50 of 24 µM (Fig. 2G), were almost completely eliminated while the KO cells continued to proliferate (Fig. 2F and Supplementary Movies 1 and 2). These results indicated specificity of 3BP for cells expressing MCT1.

### Optimal Encapsulation of 3BP protects from degradation by human serum

One of the major challenges of using 3BP as a chemotherapeutic agent is that it is rapidly inactivated by the serum proteins it alkylates before it leaves the circulation and enters cancer cells.^26^ We have previously shown that encapsulation of 3BP with a cyclodextrin can mitigate its toxicity, presumably by permitting the use of lower concentrations of the encapsulated drug compared to the free drug. ^28^ Therefore, we focused on optimizing the microencapsulation procedure and then examined the serum stability of the drug. To optimize microencapsulation of 3BP in cyclodextrins, we chose the β-cyclodextrin scaffold rather than alpha or gamma cyclodextrins based on structural modeling. ^28^ We first evaluated two types of β-cyclodextrin modifications, one with the hydroxyls substituted with succinyl groups and the other with the hydroxyls substituted with 2-hydroxypropyl groups. Second, for the modifications involving succinyl groups, we sought to determine the optimal number of hydroxyl substitutions. Finally, we investigated the optimal ratio of cyclodextrin to 3BP. To evaluate these various formulations, we first used size exclusion chromatography to ensure that the amount of free 3BP was minimized (Fig. 3A). Second, we used cell toxicity as a measure of the stability of these formulations. Because the cytotoxicity of 3BP decreases upon interaction with serum proteins, residual cell toxicity following exposure to serum is directly related to its resistance to serum degradation. Therefore, the formulations were incubated with human serum at 37 °C and aliquots were collected at various time points for assessment of cell toxicity in DLD-1 cells which have high expression of MCT1 and sensitivity to 3BP (IC50, 22.64 µM; Fig. 1B). Among the β-cyclodextrin formulations tested, we found that succinyl substituted cyclodextrins were capable of protecting 3BP in sera more efficiently than hydroxypropyl substituted versions. Moreover, a succinyl-substituted cyclodextrin with an average of 3.4 succinyl groups per cyclodextrin, at a molar ratio of 1.2 β-cyclodextrin per 3BP, performed best (Fig. 3A). This <u>M</u>icroEncapsulated formulation was named “ME3BP-7”. Although free 3BP lost 90% of its activity by 30 min of exposure to serum, ME3BP-7 lost < 10% of its activity in 30 min (Fig. 3B). Remarkably, even after 8 h of incubation with serum, ME3BP-7 retained > 70% of its activity (Fig. 3B). Hydroxypropyl-substituted cyclodextrin-3BP complexes were more stable than free 3BP, losing just over 50% of its activity after 30 min of exposure to serum. However, these complexes were not nearly as stable as ME3BP-7 (Fig. 3B).

**Fig. 3.**
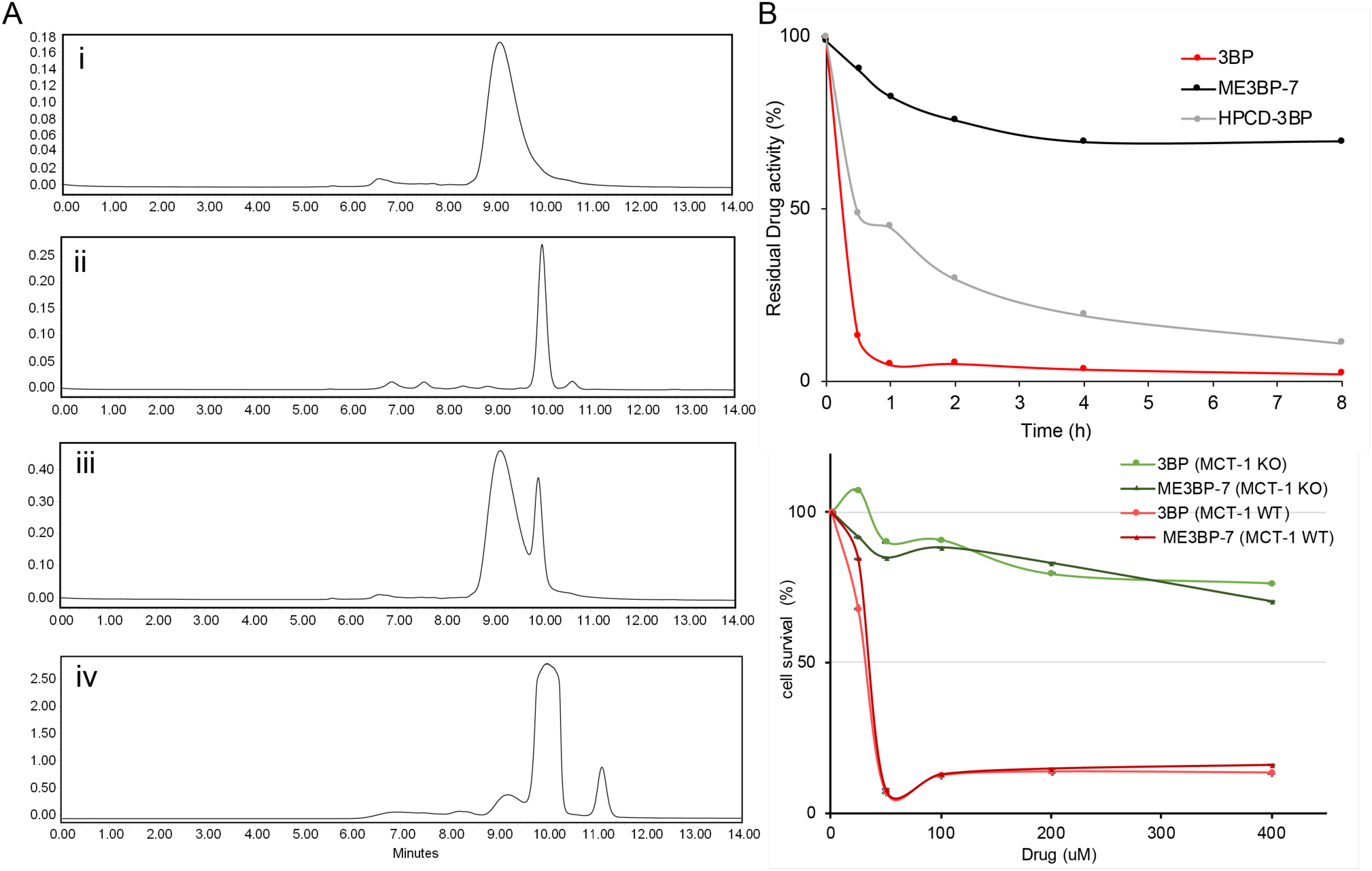
New formulations and serum stability of 3BP in cyclodextrin complexes. **(A**) HPLC: Evaluation of different microencapsulated β-cyclodextrin complexes using size-exclusion chromatography (SEC).The agents examined were i. 3BP (1 mg/mL), ii. succinyl–β-CD (20 mg/mL), iii. a mixture of 10 µL of 3BP and 10 µL succinyl–β-CD, and iv. ME3BP-7 (10 mg/mL). Samples were monitored at 220 nm as shown in A-D. (**B**) Serum stability assay using DLD-1 cells. (**C**) ME3BP-7 specificity assessed with MIA PaCa-2 parental and MIA PaCa-2 MCT-1 KO cells.

Finally, parental MIA PaCa-2 (MCT1 WT) in contrast to MCT1 KO cells were sensitive to the encapsulation of 3-BP within ME3BP-7, indicating that the compound retained specificity as well as cytotoxicity for cells expressing MCT1 (Fig. 3C).

### ME3BP-7 results in rapid target-cell killing

We also noticed that the morphology of the MIA PaCa-2 cells rapidly changed in response to exposure to 3BP or ME3BP-7. To determine the significance of this change in morphology, we performed washout experiments on co-cultures of WT and KO cells exposed to drugs. Co-cultures were exposed to 3BP or ME3BP at various concentrations and time periods, and cell proliferation was followed for 48 h after removal of the drug. Exposure to either 3BP or ME3BP-7 for as little as 30 min led to loss of the majority of WT MIA PaCa-2 cells. In contrast, KO MIA PaCa-2 cells continued to proliferate under all concentrations of the drugs and time points tested (Fig. 4A-D and Supplementary Fig. 1-2). In addition, gemcitabine or any of the FOLFIRINOX agents, including irinotecan, oxaliplatin (Supplementary Movie 3), and 5-FU, exhibited no differential effect on WT and KO cells, thus emphasizing the specificity of 3BP and ME3BP for MCT1 expressing cells. Moreover, 3BP and ME3BP-7 caused cell death of MIA PaCa-2 cells, while the standard of care agents only decreased their proliferation (Fig 4 and Supplementary Fig. 1-2).

**Fig. 4.**
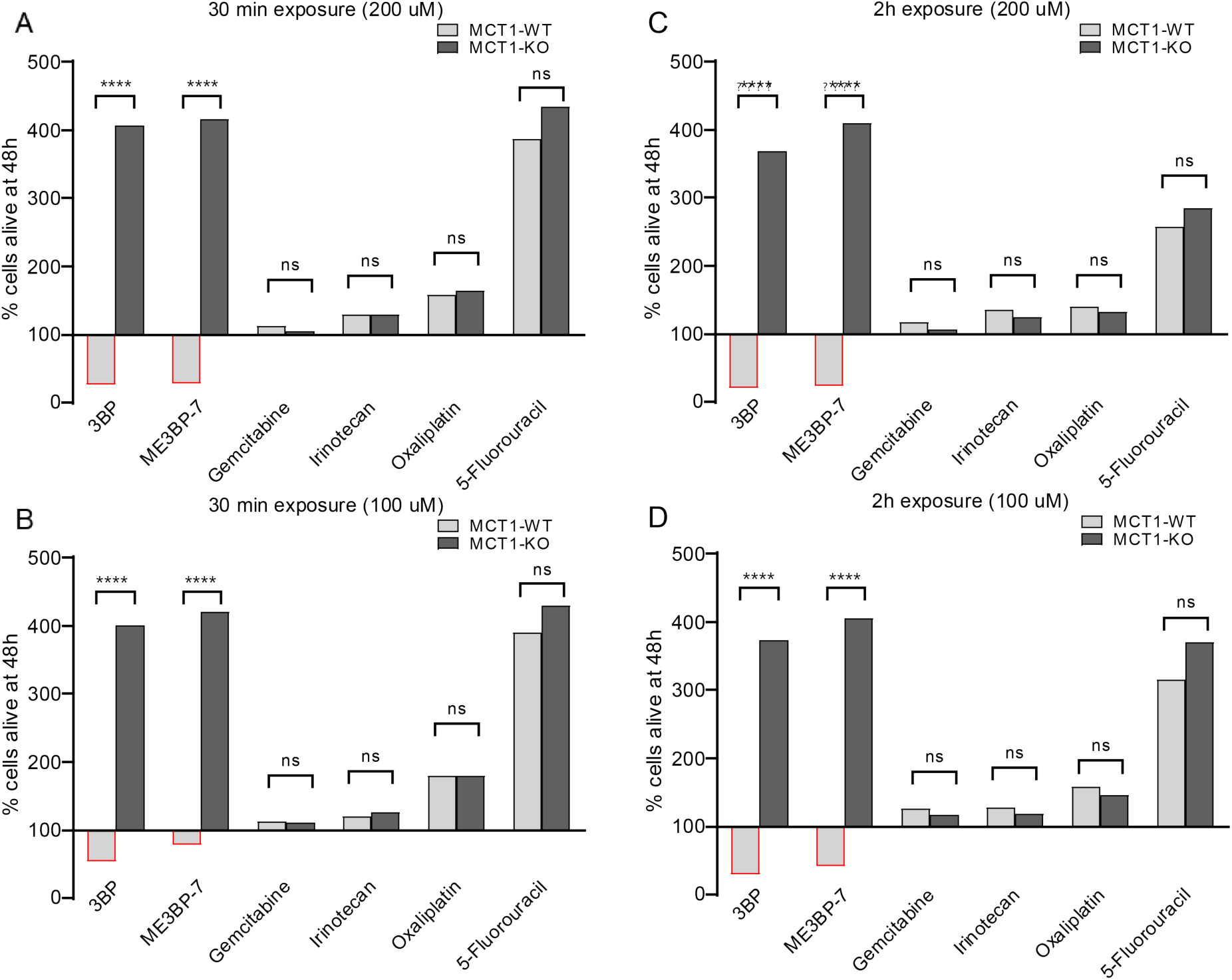
Comparison of MCT-1 specific cytotoxicity of 3BP, ME3BP-7 and current standard of care agents for PDAC upon short exposures. Viability of MCT-1 isogenic panel after (**A**) Drug exposure for 30 minutes at 200 uM. (**B**) Drug exposure for 2 hours at 200 uM (**C**) Drug exposure for 30 minutes at 100 uM (**D**) Drug exposure for 2 hours at 100 uM

### ME3BP-7 shows efficacy *in vivo*

To assess efficacy in vivo, we treated mouse models of PDAC with ME3BP7. We first tested the feasibility of delivering ME3BP-7 systemically and compared the toxicity of free 3BP with that of ME3BP-7 in athymic nude mice. Escalating doses ranging from 8.0-33 mg/kg of free 3BP (as ME3BP-7) were used to identify a maximum tolerated dose of ME3BP-7. Equivalent amounts of free 3BP and ME3BP-7 were infused via vascular access buttons (33 mg/kg of 3BP) in 200 µL of PBS every other day for one week. In preliminary experiments, two-thirds of mice in the free 3BP group became ill, showing severe weight loss, pallor and decreased activity, and died after only two doses. No overt toxicity was evident even after five weeks of treatment with ME3BP-7 at the same dose (33 mg/kg) (Supplementary Fig 3A).

We then generated orthotopic xenografts with Panc 02.13, a patient-derived PDAC cell line with high expression of MCT1 (RNA expression of 137 transcripts per million (Fig. 1B), engineered to express firefly luciferase. Tumor bearing animals were randomized into three treatment arms on the basis of intra-vital multiphoton imaging (IVI) and treatment was initiated a day later (Fig. 5A). Guided by our dose finding toxicity studies, no treatment arm with free 3BP was included in this study as none of the animals would survive the course of the therapy. Over the course of four weeks of treatment (dosed Mondays, Wednesdays, and Fridays), no discernable adverse effects or weight loss was observed in the treated animals (Supplementary Fig 3A). Animals in both treatment cohorts showed near complete abrogation of tumor growth (p<0.01, 1-way ANOVA, Fig. 5B and 5C). Finally, the weight of residual tumors after four weeks of ME3BP-7 was significantly decreased in treated animals relative to controls (p=0.01, Mann Whitney U Test, Fig. 5D).

**Fig. 5.**
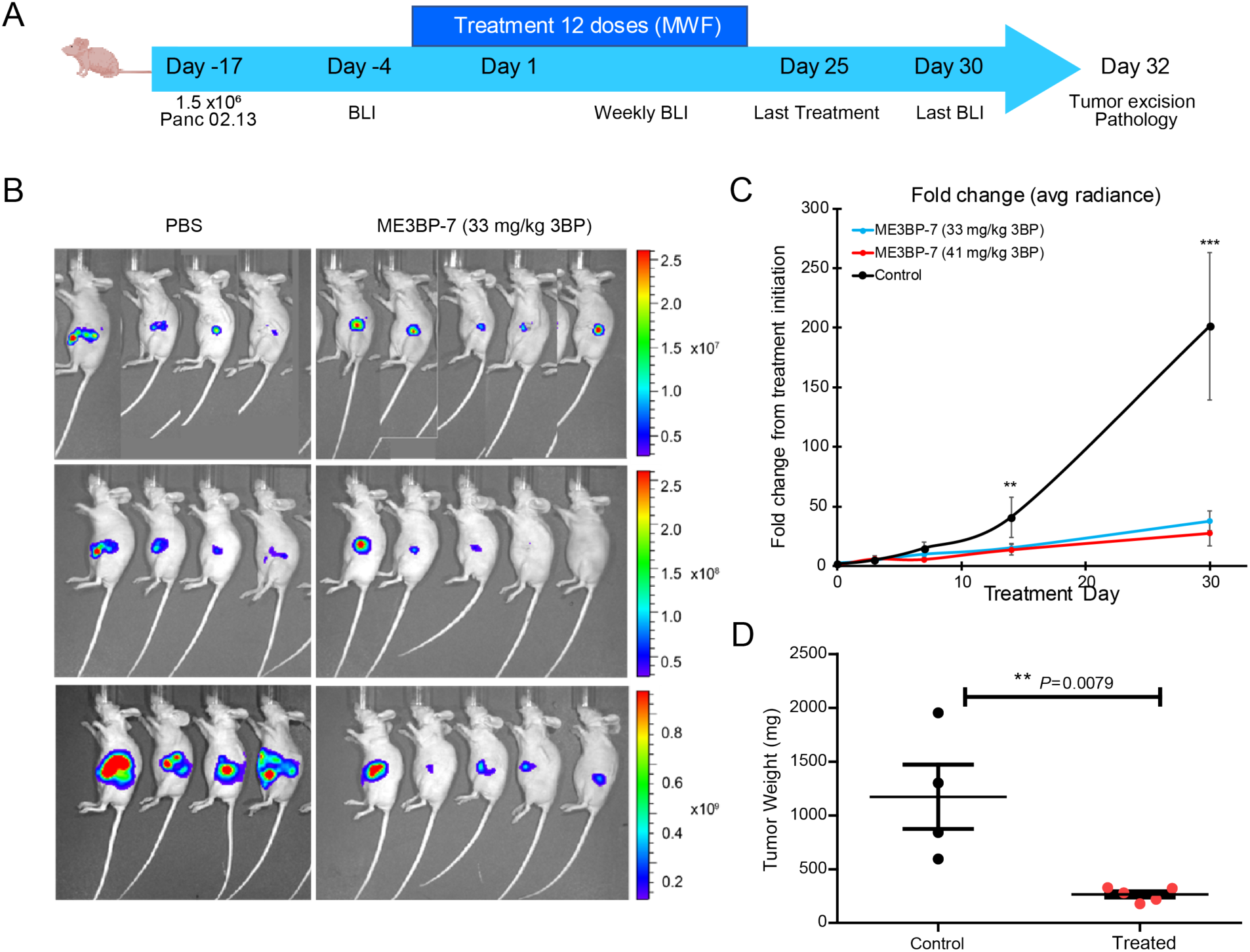
ME3BP-7 inhibits tumor growth of orthotopically implanted pancreatic cancer cell line Panc 02.13 with high MCT1 expression. (**A**) Timeline and design of in vivo tumor experiments. (**B**) Bioluminescence images of nude mice bearing orthotopic Panc 02.13 tumors. (**C)** Mean fold change in radiance from day of treatment initiation (** *P* < 0.01, *** *P*<0.001 1 way ANOVA. (**D**) Weights of residual tumors harvested upon termination of therapy (**\*\*** *P* < 0.01 Mann-Whitney U test).

We next tested the efficacy of ME3BP-7 in an orthotopic xenograft model generated with patient-derived PDAC TM01212 tumor chunks. Orthotopic TM0212 xenografts expressed moderately high levels of both MCT1 RNA (37 transcripts per million) and protein (Fig. 6B). Histological evaluation revealed the development of duct-like structures within these xenografts (Fig. 6B). More importantly, all control animals developed metastases to the lung and liver, mimicking the behavior of human PDACs (Fig. 6F-H). However, only 3/7 mice treated with ME3BP-7 had any distant metastases, and the number and size of metastases in the treated vs. untreated mice were significantly decreased in treated animals relative to controls (p<0.01, Mann Whitney U Test, Fig. 6H). Again, histological evaluation of major organs did not reveal any pathological changes of significance as a result of potential toxicity in the treated mice (Supplementary Fig. 4 and Supplementary Table 1).

**Fig 6.**
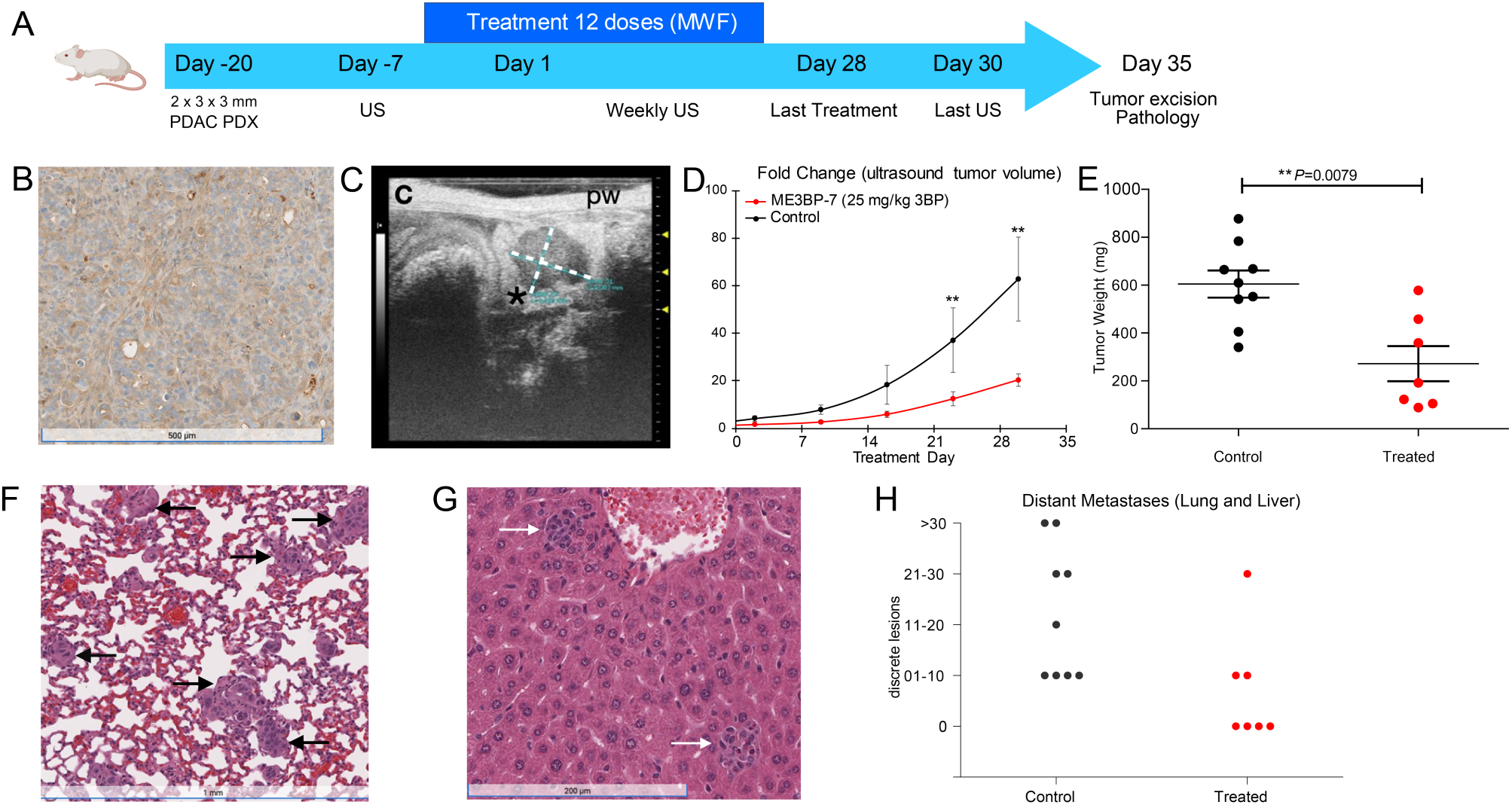
ME3BP-7 reduces tumor burden in orthotopically implanted human patient derived xenograft TM01212 with diffuse expression of MCT1. **A**) Timeline and design of in vivo therapeutic study. **B**) IHC of orthotopic PDx TM01212 showing diffuse but uniform expression of MCT1. (**C**) Representative ultrasound image of orthotopically implanted tumors in NSG mice. (**D**) Mean fold change in tumor volume (n = 10) from day of treatment initiation. (**E**) Weights of residual tumors harvested upon termination of therapy. (**F** and **G**) H&E of lung and liver with metastases from untreated animals. (**H**) Number of metastatic lesions harvested from control and treated mice upon termination of therapy. (**\*\*** *P* < 0.01 Mann-Whitney U test)

We evaluated efficacy of ME3BP-7 in a second PDAC patient-derived xenograft model, TM01098 (Supplementary Fig 5A). Although these tumors cells expressed high levels of *MCT1* RNA (96 transcripts per million), protein expression exhibited focal staining (Supplementary Fig 5B), which was in contrast to the uniform expression in Panc 02.13 (Fig. 1C) and TM01212 xenografts (Fig. 6B). Despite focal MCT1 protein expression, significantly slower growth of TM01098 xenografts occurred in animals treated with ME3BP-7 compared to controls, based on US (p<0.01, Mann Whitney U Test, Supplementary Fig. 5D) and tumor weight (p=0.0028, Supplementary Fig. 5E).

### Analysis of expression of MCT1 in human tissues

To further validate the mRNA expression data in human tumors in publicly available databases ^33,34^ (Supplementary Fig. 6), immunohistochemistry was performed on a total of 95 PDACs on tissue arrays. Our analyses showed widespread MCT1 expression relative to adjacent normal cells in 30% of the cases, focal regions of high expression in 5-8%, and diffusely low or absent MCT1 expression in 60-65% (Fig. 7A and C).

**Fig. 7.**
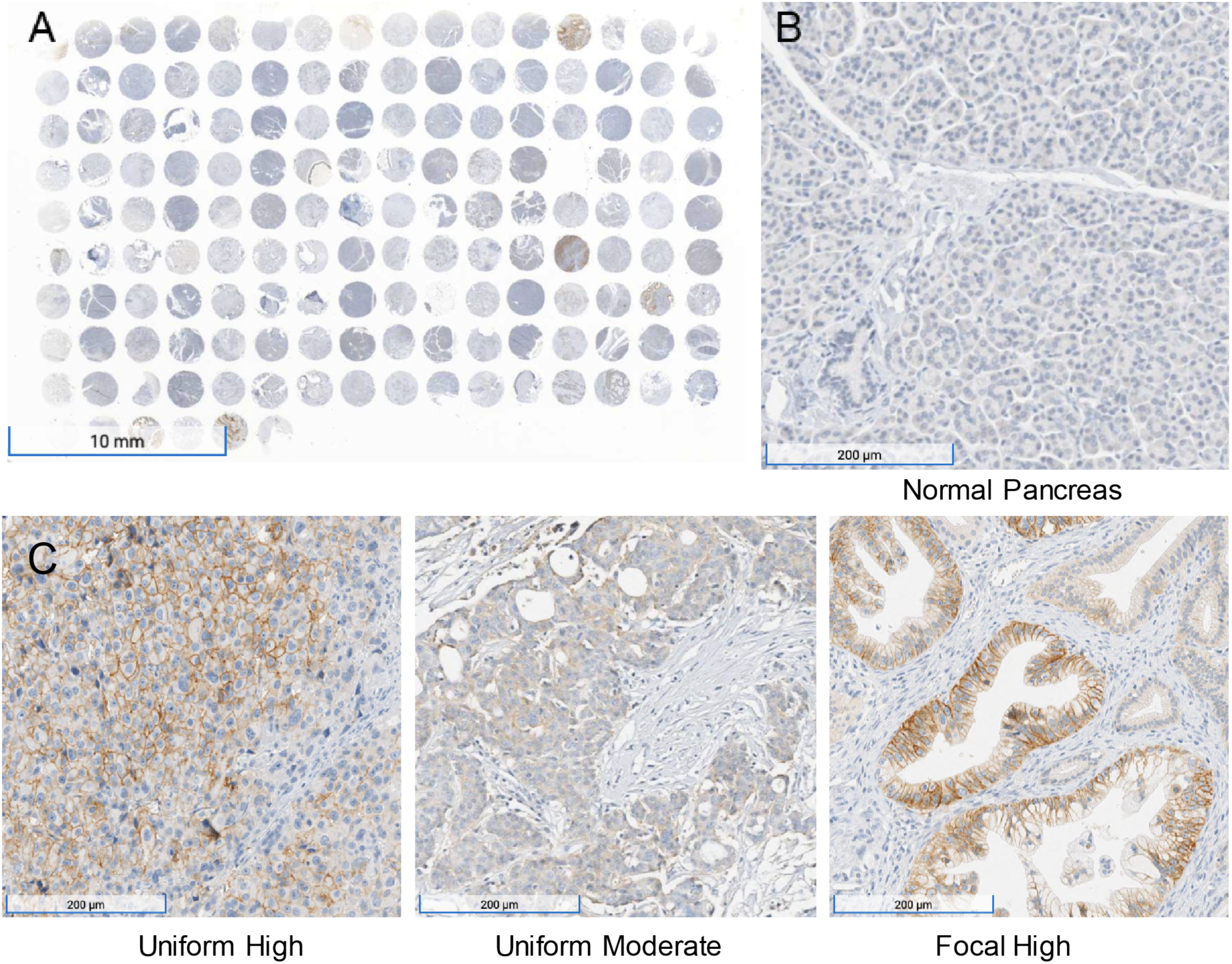
Immunohistochemistry performed on a PDAC tissue microarray with MCT1 antibody. (**A**) Overview (1x) of IHC performed on tissue microarray (HPanA150CS03, BioMax U.S) of pancreatic carcinoma cases with MCT1 antibody. (**B**) IHC of normal pancreas (10x) from the same microarray with MCT1 antibody. (**C**) Representative examples of uniform high, uniform moderate and focal high expression of MCT1 in human PDAC samples from the array (10x).

None of the 51 normal tissues recorded in the GTEX database (https://gtexportal.org/home/gene/SLC16A1) expressed average levels of *MCT1* RNA greater than the approximate threshold required for sensitivity to 3BP in cancer cell lines (∼ 50 transcripts per million, Supplementary Fig. 7). The highest expression levels in normal tissues, were found in the colon, testis, and uterus (median 37, 38, and 32 transcripts per million, respectively, among 142 to 376 tissue samples assessed for each tissue type). ^35^ The RNA levels were consistent with the immunohistochemistry; staining was low or absent in normal pancreas, brain or ovary while readily observed in testis, colon and uterine tissue (Supplementary Fig. 8). On the basis of this information, we carefully examined ME3BP-7 treated mice at necropsy and found no discernible damage after 4 weeks of treatment in either the colon or uterus, while the pancreatic cancers showed extensive necrosis (Supplementary Table 1 and Supplementary Fig. 4).

## DISCUSSION

Pancreatic cancer is a major cause of cancer morbidity and mortality, and is predicted to be the third leading cancer killer in the next decade. Average life expectancy is under a year in advanced disease and there is an undeniable unmet need for better therapeutics. Public datasets reveal that sizable fraction of pancreatic cancers express relatively high levels of a membrane protein, MCT1, ^33^ while the same is barely detectable in normal pancreas. Despite the promise, specific small molecule inhibitors of MCT1 have not shown any benefit in human clinical trials, rendering MCT1 an unrealized but exciting biological target.

The results described in this study offer an innovative therapeutic approach using ME3BP-7 to target MCT1. The closest successful precedents for ME3BP-7 are antibody-drug conjugates such as trastuzumab emtansine^36^ or inotuzumab ozogamicin.^37^ ME3BP-7 differs from antibody-drug conjugates in several important respects. First, ME3BP-7 is a small molecule, drastically reducing drug production challenges and cost. Second, ME3BP-7 itself mediates the targeting as well as the toxicity, while antibody-drug conjugates use an antibody for targeting and a separate, small molecule to kill cells.^38–40^

One of the most challenging aspects of the current study was delivery of ME3BP-7 to the mice. After multiple attempts to deliver various 3BP formulations through other routes, we were only able to reliably deliver multiple doses of the drug intravenously, and the number of injections and time periods over which we could administer the drug were limited. Importantly, this logistical issue is peculiar to small animals such as mice, in which repeated vascular access is difficult. In humans, delivery of anti-cancer drugs through pump-driven intravenous or intra-arterial catheters is routinely practiced.^41–43^

Although we focused on pancreatic cancers in this study, MCT1 is overexpressed in subsets of mesotheliomas, leukemias, and several other tumor types (https://www.proteinatlas.org/ENSG00000155380-SLC16A1/pathology). As with any targeted agent, we expect one important mechanism of resistance will be loss of the target. ^44–46^ With targets that are the products of oncogenes, simple loss will not suffice, as the oncogene protein product is necessary for cell proliferation. In such cases, mutations in other sites of the protein that make it resistant to the drug, or other genes in the same pathway, generally occur. ^47–50^ MCT1 is not an oncogene, so loss of MCT1 is conceivable.

The results in Fig. 3 show that ME3BP-7 is uniquely fast-acting compared to other drugs now used in the clinic to treat patients with pancreatic cancer. Even brief exposure to ME3BP-7 results in effective cell killing of the majority of cells. On the other hand, a limitation of our study is that, despite these striking effects in vitro, we were not able to induce true regressions of extant tumors in mice, though growth was slowed (Fig. 5, 6 and Supplementary Fig. 5) and metastases were markedly reduced (Fig. 6H). We speculate that different methods for administering ME3BP-7 may improve its in vivo potency. In addition, potentially synergistic combinations with new and existing agents could also be explored to augment the potency as well as broaden the therapeutic potential of ME3BP-7. Our data suggests that combination of Irinotecan, or liposomal irinotecan as described in the NAPOLI-3 trial^51,52^, with ME3BP-7 could improve the potency as well address the pockets of heterogeneity with low MCT1in pancreatic cancer (Fig. 4 and Supplementary Fig. 1). Similarly, ME3BP-7 could alleviate resistance to glutaminase inhibition leading to an efficacious combination^53^. Finally, severe extracellular acidity around solid tumors, due to MCT1 overexpression causing immune exclusion^54^, suggests a combination of immune checkpoint inhibitors with ME3BP-7 could improve therapeutic outcomes. However, given the limitations of administering the drug to small animals, we believe that clinical trials will be the only way to reliably determine whether ME3BP-7 could serve as an adjuvant therapy for patients with pancreatic cancers. Our results highlight that ME3BP-7 shows efficacy across multiple pancreatic cancer model systems and potentially represents a powerful new therapeutic agent for this cancer type and perhaps others that express the drug’s target protein.

## MATERIALS AND METHODS

### Study Design

We designed a new formulation of the cytotoxic agent 3BP to be resistant to serum degradation and systemically administered. Modified cyclodextrins were investigated for their efficiency of encapsulating 3BP, and activity of the encapsulated drug was analyzed after exposure to serum. We used CRISPR-mediated technologies to knock out the cell membrane receptor for 3BP, MCT1, in pancreatic cancer cell lines in vitro. We also modified pancreatic cancer cell lines to express fluorescent or bioluminescent markers for the purposes of tracking cellular responses to ME3BP-7 as well as other drugs commonly used in the clinic for treating PDAC patients. Tumor cells or tumor fragments were orthotopically implanted into the pancreas of mice for in vivo experiments. Implantation was confirmed with luciferase expression in cell lines or ultrasound, followed by stratification and randomization into treatment and control groups. The MCT1-dependent killing potential of ME3BP-7 was measured with time-lapse recordings obtained with an IncuCyte® Live Cell-Analysis System (Essen Bioscience; Ann Arbor, MI, USA). Necropsies were performed for tumor resection and organ evaluation at the end of the experiment. Sample sizes for animal experiments were selected based on previous experience with the animal models but were not predetermined by power analysis. No animals were excluded from the study due to illness unless indicated. The number of replicates in each experiment is noted in the figure legends.

### Ethics statement

Six- to eight-week-old female mice were maintained according to the JHU Animal Care and Use Committee (approved research protocol MO18M79).

### Synthesis of encapsulated 3BP

Drug batches were generated as previously described. In brief, a solid sample of 3BP (1 mmol, 167 mg) was added in small portions to a solution of the corresponding cyclodextrin (1.0-1.2 mmol) in 25 mL of deionized (DI) water under constant agitation. After complete addition over 15-20 min, the samples were further agitated for 1-4 h at 25 °C. Finally, the sample was flash-frozen in liquid nitrogen or dry ice/acetone and lyophilized overnight. The lyophilized reagent was diluted in PBS 15 and 30 min prior to in vitro and in vivo experiments, respectively.

### Size-exclusion HPLC chromatography

SEC chromatography was performed with a Shodex-OH Pak at an elution rate of 1 mL/min of PBS under isocratic conditions and monitored at 220 nm. The samples included i) free 3BP (1 mg/mL); ii) succinyl–β-CD without 3BP (20 mg/mL); iii) a mixture of 10 µL of free 3BP at 1 mg/mL and 10 µL ME3BP-7 at 10 mg/mL; and iv) ME3BP-7 (10 mg/mL).

### Cell culture

Human PDAC cell lines, including MIA PaCa-2, PSN-1, Panc 02.13, AsPC-1, BxPC-3, and CFPAC-1, were obtained from the American Type Cell Culture (ATCC) (Manassas, VA, USA). Cell lines were maintained at 37 °C in a humidified 5% CO2 atmosphere in T75 tissue culture flasks containing 14 mL of the medium recommended by ATCC (DMEM for MIA PaCa-1 parental and MCT1-KO; RPMI 1640 for Panc 02.13, AsPC-1, BxPC-3, and DLD-1; and IMDM for CFPAC-1. All media were supplemented with 10% fetal bovine serum (Mediatech, Inc.; Manassas, VA, USA) and 1 % penicillin-streptomycin. Cells were passaged at the ATCC recommended ratio every 4-5 days with trypsinization and resuspended in fresh medium in a new flask. Only early passaged cells harvested at 70-85% confluence were used for *in vitro* and *in vivo* experiments.

### Lentiviral transduction

MIA PaCa-2 or Panc 02.13 cells were transduced with a CMV-Firefly luciferase lentivirus carrying a puromycin-selectable marker (Cellomics Tech; Halethorpe, MD, USA) according to the manufacturer’s instructions. Parental and MCT1-KO MIA PaCa-2 cells were transduced with Lentiviral NucLight Red and Green vectors (Sartorius), respectively, carrying a puromycin-selectable marker. All transduced cells were selected in growth medium containing puromycin (4 μg/mL) for 96 h.

### Viability assays

Cells (3-12 x 10^3^) were seeded into the wells of 96-well plates and exposed to drug 24 h after plating. For MIA PaCa-2 parental and -MCT1-KO co-culture experiments, cells were suspended in medium at equal concentration and seeded at 12,000/genotype into wells. Cells in experiments were imaged with time-lapse fluorography using the IncuCyte® Live Cell-Analysis System (Essen Bioscience; Ann Arbor, MI, USA). For green- and red-labeled cells, a processing definition for the IncuCyte®was was created, and the green or red fluorescence channels were used for analysis. All *in vitro* experiments were conducted at least in duplicate. Green and red object counts were used to track parental and MCT1-KO MIA PaCa-2 experiments, while confluence was used to track unlabeled cells. Cell viabilities were recorded as the fraction of cells that survived treatment at the indicated drug molarities relative to the control wells. Dose-response curves were fit using Prism 9 software.

### Serum stability assays

Free 3BP, ME3BP-7, and HPCD-3BP were incubated at various concentrations (12.5 μM, 25 μM, 50 μM, 100 μM, and 200 μM) with 90 μL of human serum (Sigma, Cat No. H3667; St. Louis, MO, USA) at 37 °C for up to 8 h. Aliquots were collected at 30 min, 1 h, 2 h, 4 h, and 8 h and stored at −80°C until further analysis. For the cell toxicity assay used to assess the amount of biologically active 3BP at each time point, parental DLD-1 (MCT1, 387 TPM) cells were plated at 35-40% confluence, and exposed to 10 µL of the collected drug sample diluted with 190 µL of medium. Cell death was measured by imaging cells after 72 h in culture with the IncuCyte® Live Cell-Analysis System (Essen Bioscience). Residual drug activity was estimated by comparison of cell death induced by drug samples incubated with human serum for different periods of time, to the cell death induced by an unincubated drug sample to derive the % drug activity.

### Timed exposures to the drug

Cells were seeded into 96-well plates at a density of 24,000 cells per well (12,000 parental MIA PaCa-2 red and 12,000 MCT1-KO MIA PaCa-2 green). After 24 h, cells were treated with vehicle (complete DMEM media) or a serial dilution of a drug (bromopyruvic acid, Sigma-Aldrich; gemcitabine HCl, irinotecan, oxaliplatin, and 5-fluorouracil, Selleck Chemicals; Houston, TX, USA). Each treatment condition was conducted in triplicate unless otherwise stated. The drugs were either removed from the plates at 30 min or 2 h, or left on cells. Wells were gently washed with 200 µL of PBS twice and replaced with fresh medium. Residual cells were assessed with IncuCyte® Live Cell-Analysis System. Cell viabilities were reported as the percentage of cells remaining after treatment at the indicated drug molarities relative to the average of three control wells without drug.

### Genetic inactivation of *SLC16A1* (MCT1)

The Alt-R CRISPR system (Integrated DNA Technologies, IDT; Coralville, IA, USA) was used to delete *SLC16A1*, the gene encoding MCT1 protein, in the DLD-1 and MIA PaCa-2 cell lines. The gRNA sequence was designed with CHOPCHOP v3 ^55^. Alt-R CRISPR Cas9 crRNAs (ACCATGCCATTCAGGCTAGT, IDT; SEQ ID NO:1) and Alt-R CRISPR-Cas9 tracrRNA (1072532, IDT) were resuspended in Nuclease-Free Duplex Buffer (IDT) at a concentration of 100 μM. The crRNAs and tracrRNA were mixed in a 1:1 molar ratio and denatured for 5 min at 95 °C, followed by slow cooling to room temperature for duplexing. Cas9 Nuclease (1081059, IDT) was then added at a 1.2:1 molar ratio and allowed to stand at room temperature for 15 min. Forty pmoles of the Cas9 ribonucleoprotein (RNP) containing tracrRNA/MCT1 crRNA duplex was mixed with 2 x 10^5^ cells in 20 μL of OptiMEM (31985088, ThermoFisher Scientific; Waltham, MA, USA), loaded into a 0.1 cm cuvette (1652089, Bio Rad; Hercules, CA, USA) and electroporated at 120 V for 16 ms using an ECM 2001 (Harvard Apparatus; Holliston, MA,

USA). Cells were immediately transferred to complete growth medium and cultured for one week. Upon reaching confluence, cells were plated at a density of 0.5 to 2 cells per well in 96-well plates and cultured for 3 weeks. Single colonies were transferred into 2 replica 96-well plates. Genomic DNA was harvested from one of the plates using the Quick-DNA™ 96 Kit (Zymo Research; Orange, CA, USA) and PCR amplified using Q5® Hot Start High-Fidelity 2X Master Mix (New England BioLabs; Ipswich, MA, USA). SafeSeqS^56^ was used to confirm the mutation status of selected clones. Single MIA PaCa-2 clones deleted for *SLC16A* were collected from the matched replica plate, and six clones were pooled at equal ratios for further experiments.

### MCT1 immunostaining

DLD-1 parental and isogenic DLD-1-MCT1 KO cells were used as the positive and negative controls for testing the specificity of five commercially available MCT1 antibodies. Cells were trypsinized, washed in media to inactivate the trypsin, pelleted at 400x*g* for 10 min, fixed in 10% formalin and embedded in paraffin. Sections (4 µm) were cut from the blocks and incubated with different dilutions of primary and species-appropriate secondary antibodies. The anti-MCT1 mouse antibody from Santa Cruz Biotechnology (SC-365501, Lot number D2319; Dallas, TX, USA) was chosen for immunostaining with a Ventana Discovery Ultra autostainer (Roche Diagnostics; Indianapolis, IN, USA) at the Oncology Tissue Services Core of Johns Hopkins University School of Medicine. Briefly, following deparaffinization and rehydration of sections, epitope retrieval was performed in Ventana Ultra CC1 buffer (catalog# 6414575001, Roche Diagnostics) at 96 °C for 64 min. Antibodies were diluted in antibody dilution buffer (catalog# 5280524001, Roche Diagnostics; anti-MCT1 antibody, 1:2000 for cell pellets, 1:200 for patient derived xenografts, and 1:100 for other tissues or tissue microarrays) and incubated with the slides at 36 °C for 60 min. Following standard washing in the Ventana auto-stainer, bound antibodies were detected with an anti-mouse HQ detection system (catalog# 7017936001 with 7017782001, Roche Diagnostics) and a Chromomap DAB IHC detection kit (catalog # 5266645001, Roche Diagnostics). The slides were then counterstained with Mayer’s hematoxylin, dehydrated, and mounted in Toluene mounting medium (MER 7720, Mercedes Scientific; Lakewood Ranch, FL, USA).

### Tissue microarrays

IHC for MCT1 was performed on two commercially available FFPE tissue microarrays (BC001130 and HPanA150CS03; US Biomax, Inc.; Rockville, MD) for a total of 100 cases of pancreatic cancer. BC001130 included 20 cases of pancreatic carcinoma (triplicate cores per case), and HPanA150CS03 included 80 cases with adjacent normal tissue. Two independent reviewers, including a pathologist, scored the intensity and pattern of MCT1 staining for tumor and normal tissues. Any differences in grading were reviewed and resolved. Upon evaluation, 5 samples were considered to be of poor quality and disregarded. Of the 95 samples analyzed, 56% (n = 53) were positive for MCT1 and 44% (n = 42) were negative. The staining intensity of the 53 MCT1 positive pancreatic cancer tissues was scored as follows: 58% (n = 31), low intensity; 30% (n = 16), intermediate intensity; and 12% (n = 6) high intensity.

### Animal experiments

Escalating doses ranging from 8.0-33 mg/kg of free 3BP (as ME3BP-7) were infused via the tail vein of healthy nude mice to determine the maximum tolerated dose. Repeated tail vein injections of free 3BP caused significant scarring of the tail, limiting use for sustained injections of the free drug, while injections of ME3BP-7 exhibited minimal sclerosis. For optimal comparison with the free drug, implantable vascular access buttons (VAB) were used to provide direct access to the jugular vein for accurate administration of both drugs. Crl:NU(NCr)-*Foxn1^nu^* (athymic nude mice; Charles River (490); Wilmington, MA, USA; Fig. 5) were used for experiments performed with pancreatic tumor cell lines. More severely immunodeficient mice were used for patient-derived xenografts which did not uniformly grow in athymic nude mice. In addition, different types of severely immunocompromised mice were used for the patient-derived xenografts due to limited availability of animals during the COVID pandemic: NOD.Cg-Prkdc Il2rg /SzJ ((NSG), the Jackson Laboratory (005557); Bar Harbor, ME; Fig. 6); and NOD-*Prkdc^em26Cd52^Il2rg^em26Cd22^*/NjuCrl (NCG) (Charles River (572)); Supplementary Fig. 5).

#### Models

##### Panc02.13

Orthotopic tumors were generated by implanting 1.5 x 10^6^ luciferase-expressing Panc 02.13 cells (with 10% Matrigel) in the pancreas of 20 nude mice. On day 13 post-implantation, tumor burden was assessed with the IVIS Live cell imaging system, and 15 animals with similar bioluminescence signals were stratified and randomized into 3 cohorts. On day 16, a second baseline image was recorded, and treatment was initiated on day 17 post-implantation. Animals in the control group (n = 4) were administered 200 µL of PBS (vehicle), while the two treatment arms received 33 (n = 5) and 41 mg/kg (n = 4) of 3BP in the form of ME3BP-7 in 200 µL of PBS. Body weights were measured immediately before drug administration to ensure appropriate dosage and delivery. Treatments were administered by a single intravenous (i.v.) bolus delivered over 30 seconds for 4 weeks on each Monday, Wednesday, and Friday (total of 12 injections), and bioluminescence was recorded once each week. At the end of four weeks, all animals were euthanized and their tumors weighed.

##### TM01212

Subcutaneous TM01212 xenografts were harvested from 3 hosts, and tumor pieces (2 mm x 1 mm x 1 mm) were orthotopically transferred into the pancreas of 20 NSG mice. On day 13, tumor bearing animals (take rate 90%) were identified with US and randomized into 2 groups of 10 mice each. The control group was infused with the inactive agent (270 mg/kg sCD), while the treatment group was administered ME3BP-7 formulated at 25 mg/kg of 3BP. Bolus injections via the tail vein were administered every Monday, Wednesday, and Friday for four weeks (12 total doses) starting at day 20 post-tumor implantation. Tumor growth was tracked with weekly US measurements, and animals were euthanized on day 35 after the initiation of treatment to assess tumor burden in the pancreas, lung, and liver. Two of the treated mice received < 80% of the intended dose of ME3BP-7 (due to tail vein sclerosis) and were removed from the analysis. Some mice did not receive the full dose due to tail-scarring and responded less well than the other mice.

##### TM01098

Subcutaneous TM01098 xenografts were harvested from 3 hosts, and tumor pieces (2 mm x 3 mm x 3 mm) were transplanted into 29 NCG mice (17 with surgically implanted vascular access buttons and 12 without). On day 9, tumor bearing animals were identified with US (take rate 76%). Five mice died from VAB related complications and were removed from the study. Tumor bearing hosts were separated into 2 groups; the mice in the control group (n = 7) received 200 µL of PBS *via* tail vein i.v. bolus infusion (Monday, Wednesday, and Friday), while the mice in the treatment group (n = 11) received 21 mg/kg 3BP as ME3BP-7 via a VAB on the same schedule (Supplementary Fig. 5A). US measurements were taken weekly. Treatment started on post-implantation day 14, and the hosts were subjected to this regimen for 5 weeks. Mice underwent a final ultrasound on treatment day 38 and were euthanized. Tumor burden in the pancreas and at distal sites was assessed by necropsy.

#### Surgeri es

For orthotopic models, cells or small pieces of xenografts were implanted into the pancreas of each mouse in protocols modified from previously described techniques^57^. Once under anesthesia, the skin over the left abdomen was shaved and sterilized. A horizontal 1 cm incision was made over the left upper quadrant of the abdomen, right below the ribs. The underlying peritoneum was incised, and the spleen and pancreas were located. The pancreatic tissue was gently extruded from the abdominal cavity for good visualization. Either a cell aliquot was injected with a Hamilton syringe, or a piece of tumor was sutured to the tail of the pancreas with a 7-0 polypropylene suture avoiding major vessels. The implantation of the tumor fragment was standardized by size, time from extraction to implantation, and implantation technique in the tail of the pancreas. After implantation, the pancreas was gently returned to the abdominal cavity. The abdominal wall was closed with absorbable sutures, and the skin incision was closed using wound clips. Wound clips were removed once the incision site had healed, usually ∼ 10 days after surgery.

Mice were euthanized upon termination of the experiment or when major weight loss or toxicity was observed according to JHU Animal Care and Use Committee standards. Tumors and organs were harvested, weighed, placed in 10% formalin, processed, fixed in paraffin, and sectioned. Standard H&E staining was performed on sections, and an expert comparative pathologist reviewed all presented data.

#### *In vivo*-imaging for orthotopic Panc 02.13 animal model

Luminescence quantification was performed using the IVIS imaging system and Living Image software (Perkin Elmer; Waltham, MA, USA). Mice were anesthetized at 37 °C with inhaled isoflurane in an induction chamber for 5 min and received an intraperitoneal injection of luciferin (150 mL, RediJect D-Luciferin Ultra Bioluminescent Substrate, PerkinElmer, 770505). Bioluminescence images were taken 13 minutes after injection, and control fluorescence images were acquired at each screening to confirm satisfactory intraperitoneal luminescence.

#### Ultrasound protocol for orthotopic PDX animal model

A modification of an ultrasound imaging method for pancreas in mice was used.^20^ The left flanks of mice were shaved with a clipper. Mice were then injected with 2 mL of 0.9% sterile saline intraperitoneally to increase the contrast between intrabdominal organs, anesthetized with isoflurane in the induction chamber for 5 min, and placed in the lateral recumbent position with the left flank up on the imager over a heated pad. Continuous anesthesia was applied through a face cone. Pancreatic tumors were detected and measured through acquisition of a high-resolution ultrasound obtained with the VisualSonics Vevo2100 High-Resolution Ultrasound System. A 15 mm depth US window in B-mode acquisition was used on all mice. After applying gel over the area for US visualization, images were obtained in a standardized fashion. The spleen, liver, left kidney, and pancreas were identified first to confirm correct positioning.

Tumors were hypoechoic (dark/gray), while the surrounding pancreas was hyperechoic (bright/white). The tumor sutures were the most hyperechoic and frequently marked the tumor center. Trans-axial US images of the tumor were obtained by placing the ultrasound probe parallel and distal to the rib cage with the notched side of the transducer to the left (pointing anteriorly). Multiple anterior and posterior images were taken in this plane spanning the pancreas. Longitudinal US images were then obtained by placing the US probe parallel to the mouse axis with the notch side pointing towards the head of the mouse. Videos were acquired across the tumor and at the point where the maximum tumor diameter was visualized, and multiple images were saved for measurement for the longitudinal and the trans-axial views. For mice with several discrete tumors (e.g., at the peritoneal wall), the same process was repeated for each tumor. After imaging, mice were cleaned and allowed to recover completely from anesthesia before returning to their cage. To obtain measurements, files were loaded with the ultrasound software study management function. Maximal longitudinal and transversal diameters were obtained for each tumor image. To calculate tumor volumes, the formula V= a*b^2^/2, where (a) is largest and (b) is the smallest diameter within the 4 obtained measurements (2 trans-axial and 2 longitudinal).^21^

## Statistical methods

Data are presented as the mean ± SD unless otherwise specified. Statistical analyses were performed with the tests as indicated. A P value of < 0.05 was considered statistically significant unless otherwise indicated. All analyses and graph production were performed using Prism version 9.0 (GraphPad) or Microsoft Excel.

## Acknowledgements

We would like to thank Suman Paul, Sarah DiNapoli and Tushar Nichakawade for discussions and thoughtful comments. Some of the illustrations were generated with BioRender.com.

## Funding

The Virginia and D.K. Ludwig Fund for Cancer Research

Lustgarten Foundation for Pancreatic Cancer Research

The Commonwealth Fund

The Bloomberg∼Kimmel Institute for Cancer Immunotherapy

Bloomberg Philanthropies

JRT was supported by NIH Grant R25 5R25NS065729

BJM was supported by NCI grant T32 GM136577

CB was supported by NCI Grant R37 CA230400 and the Burroughs Wellcome Career Award for Medical Scientists

## Author contributions

Conceptualization: JRT, MDM, KWK, BV, SS

Methodology: JRT, MDM, BJM, NW, EW, SI, SZ, KG, SS

Investigation: JRT, MDM, GH, TA, EW, SI, IM, SS

Analysis and interpretation of data: JRT, MDM, BJM, NW, GH, TA, EW, SI, IM, CB, KWK, SZ, BV, KG, SS

Writing – original draft: JRT, SS, BV

Writing – review & editing: JRT, MDM, BJM, NW, GH, TA, EW, SI, IM, NSA, CB, KWK, SZ, BV, KG, SS

Supervision: BV, KG, SS

## Competing interests

The Johns Hopkins University has filed patent applications related to technologies described in this paper on which JRT, MDM, NP, KWK, BV, SZ & SS are listed as inventors, including: Encapsulated agents that bind to MCT1 (WO2022055748A1), Cyclodextrin compositions encapsulating a selective ATP inhibitor and uses thereof (US20190328666A1), Methods of treating liver fibrosis by administering 3-bromopyruvate (US10751306B2). BV, KWK, & NP are founders of Thrive Earlier Detection, an Exact Sciences Company. KWK & NP are consultants to Thrive Earlier Detection. BV, KWK, NP, and SZ hold equity in Exact Sciences. BV, KWK, SS, NP, and SZ are founders of, or consultants to and own equity in ManaT Bio., Haystack Oncology, Neophore, CAGE Pharma, and Personal Genome Diagnostics. NP is consultant to Vidium. BV is a consultant to and holds equity in Catalio Capital Management. SZ has a research agreement with BioMed Valley Discoveries, Inc. CB is a consultant to Depuy-Synthes, Bionaut Labs, Haystack Oncology and Galectin Therapeutics. CB is a co-founder of OrisDx and Belay Diagnostics. The companies named above, as well as other companies, have licensed previously described technologies related to the work described in this paper from Johns Hopkins University. BV, KWK, NP are inventors on some of these technologies. Licenses to these technologies are or will be associated with equity or royalty payments to the inventors as well as to Johns Hopkins University. Patent applications on the work described in this paper may be filed by Johns Hopkins University. The terms of all these arrangements are being managed by Johns Hopkins University in accordance with its conflict of interest policies.

## Notes

### Summary of Updates

high resolution images and updated discussion

