## Supplementary Figures for "ME3BP-7 is a targeted cytotoxic agent that rapidly kills pancreatic cancer cells expressing high levels of monocarboxylate transporter MCT1"

**Supplementary Figures and Movies**

Placeholder for Supplementary Movie 1 and 2

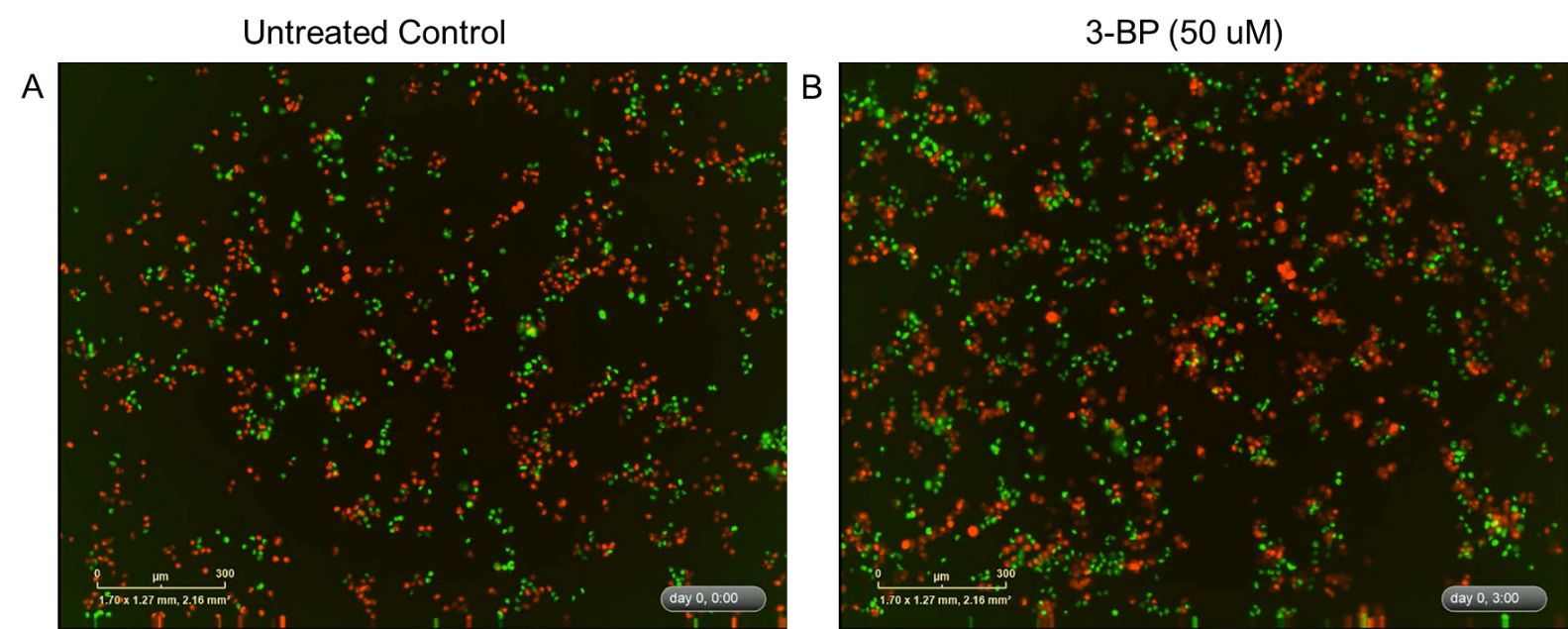

**Supplementary Movies 1 and 2:**

Time lapse movies of specific killing of MCT1 expressing cells. (**A**) Untreated controls and (**B**) with 3-BP.

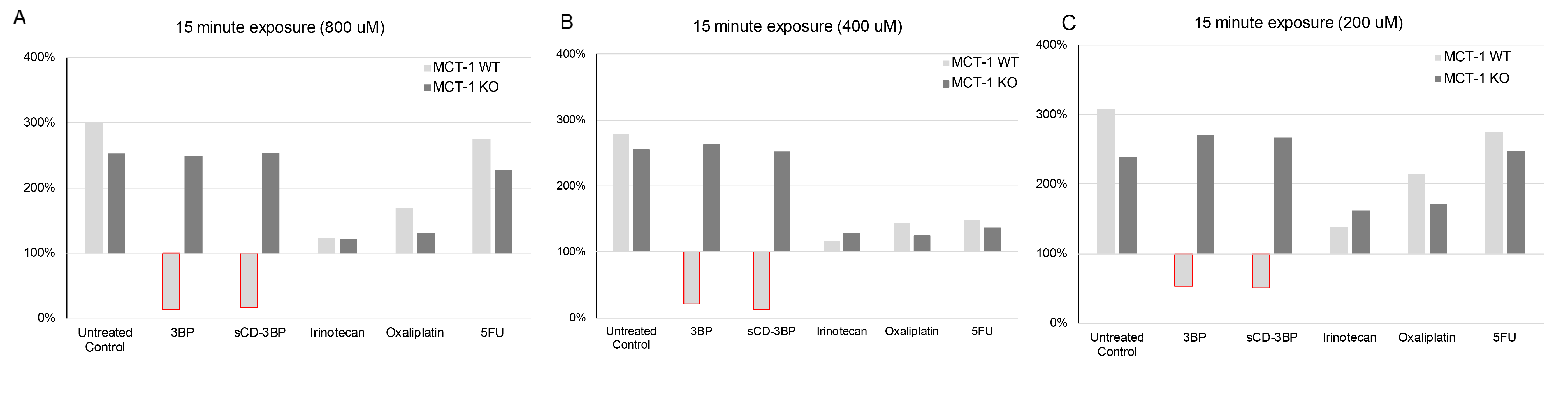

**Supplementary Fig. 1**: Incucyte data, for 15 min exposure for higher doses of various drug

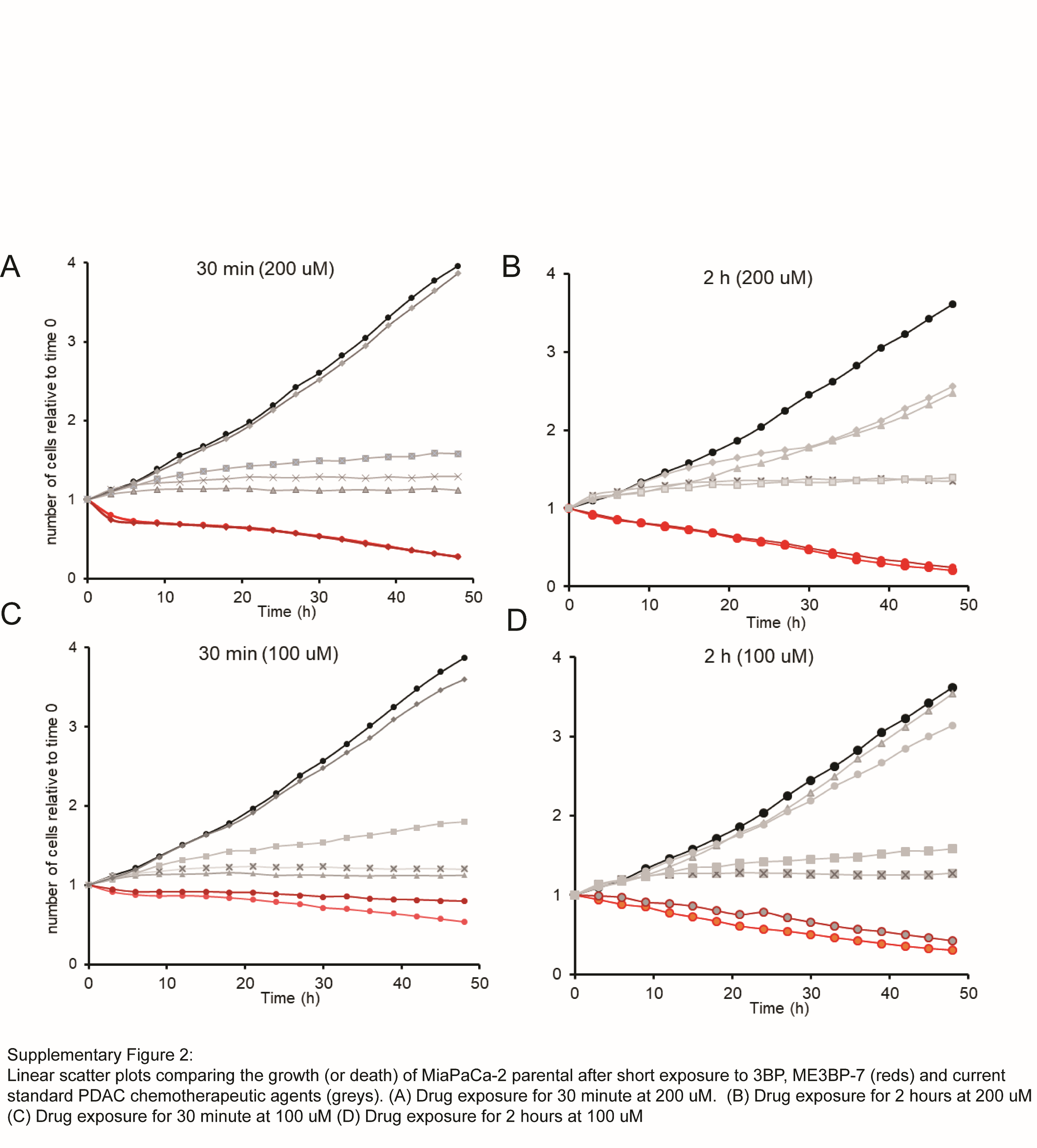

**Supplementary Figure 2.** Linear scatter plots comparing the growth (or death) of MiaPaCa-2 parental after short exposure to 3BP, ME3BP-7 (reds) and current standard PDAC chemotherapeutic agents **(**greys**).** (**A**) Drug exposure for 30 minute at 200 uM. (**B**) Drug exposure for 2 hours at 200 uM (**C**) Drug exposure for 30 minute at 100 uM (**D**) Drug exposure for 2 hours at 100 uM

Placeholder for Supplementary Movie 3

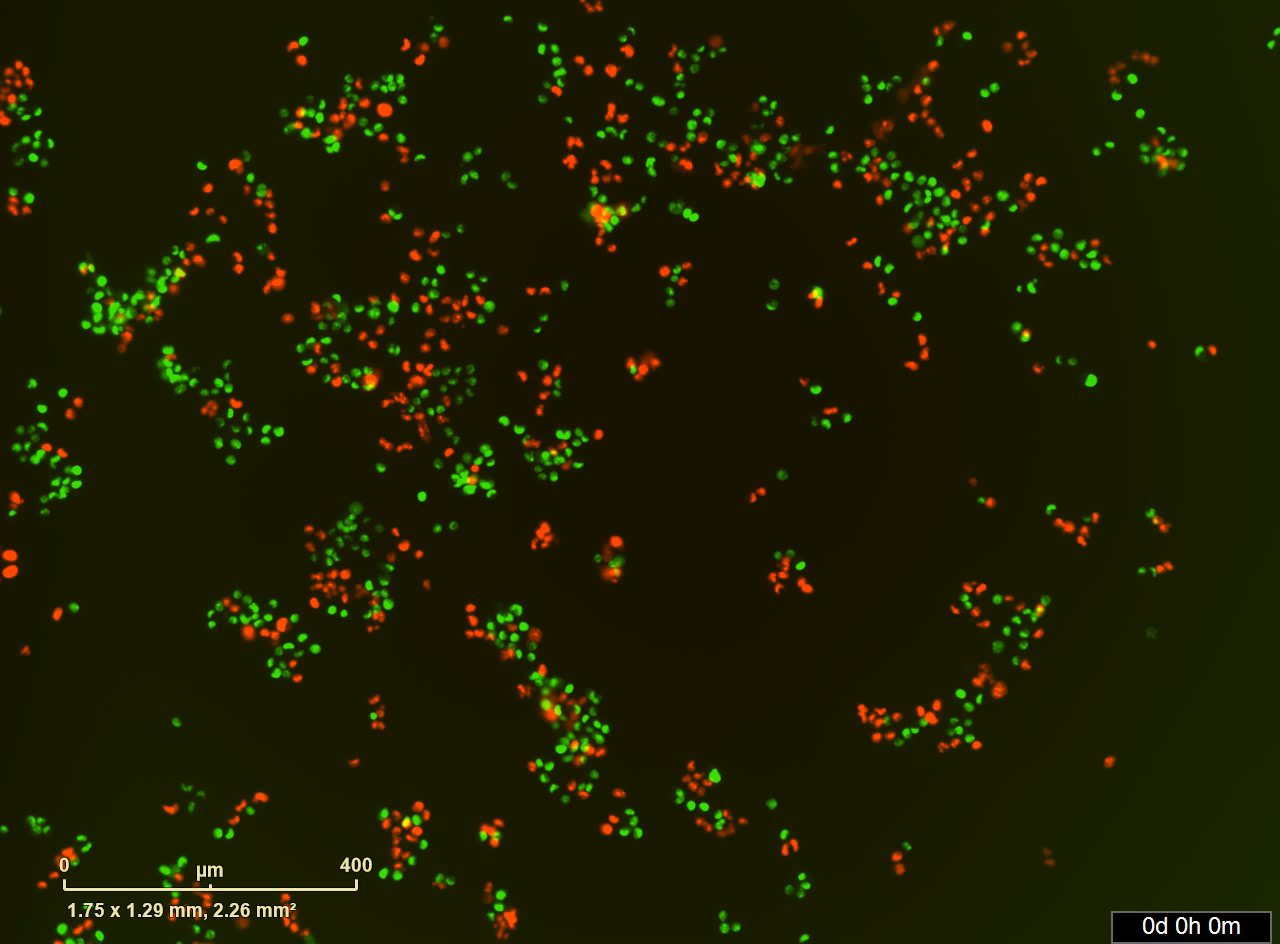

**Supplementary Movie 3**: Time lapse movie of MCT WT vs KO exposed to oxaliplatin (200 uM) for 2h.

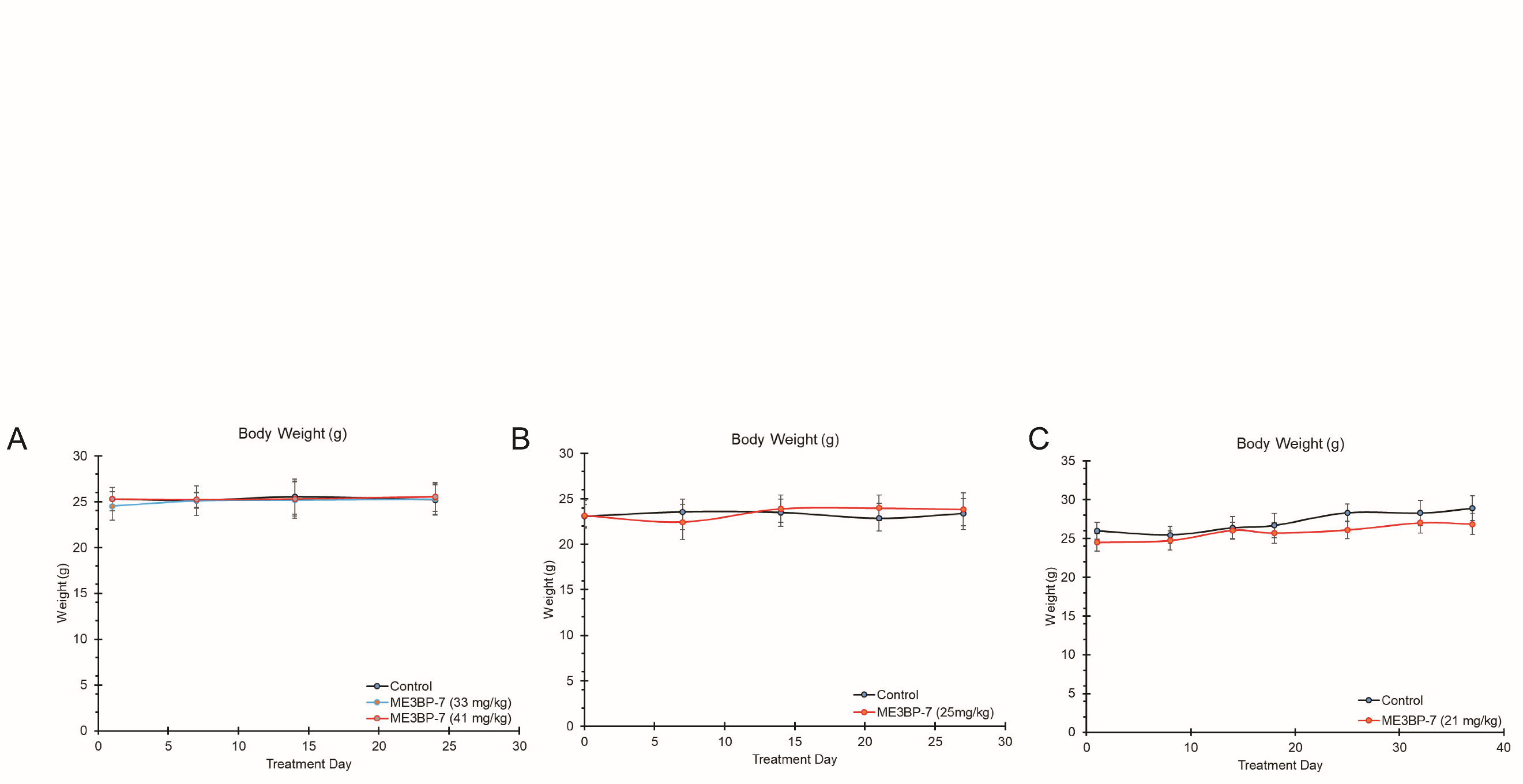

**Supplementary Figure 3**: Body weight changes over the course of ME3BP-7 administration in various murine models (**A**) Panc 02.13 in nude mice, (**B**) TM01212 in NSG mice and (**C**) TM01098 in NCG mice

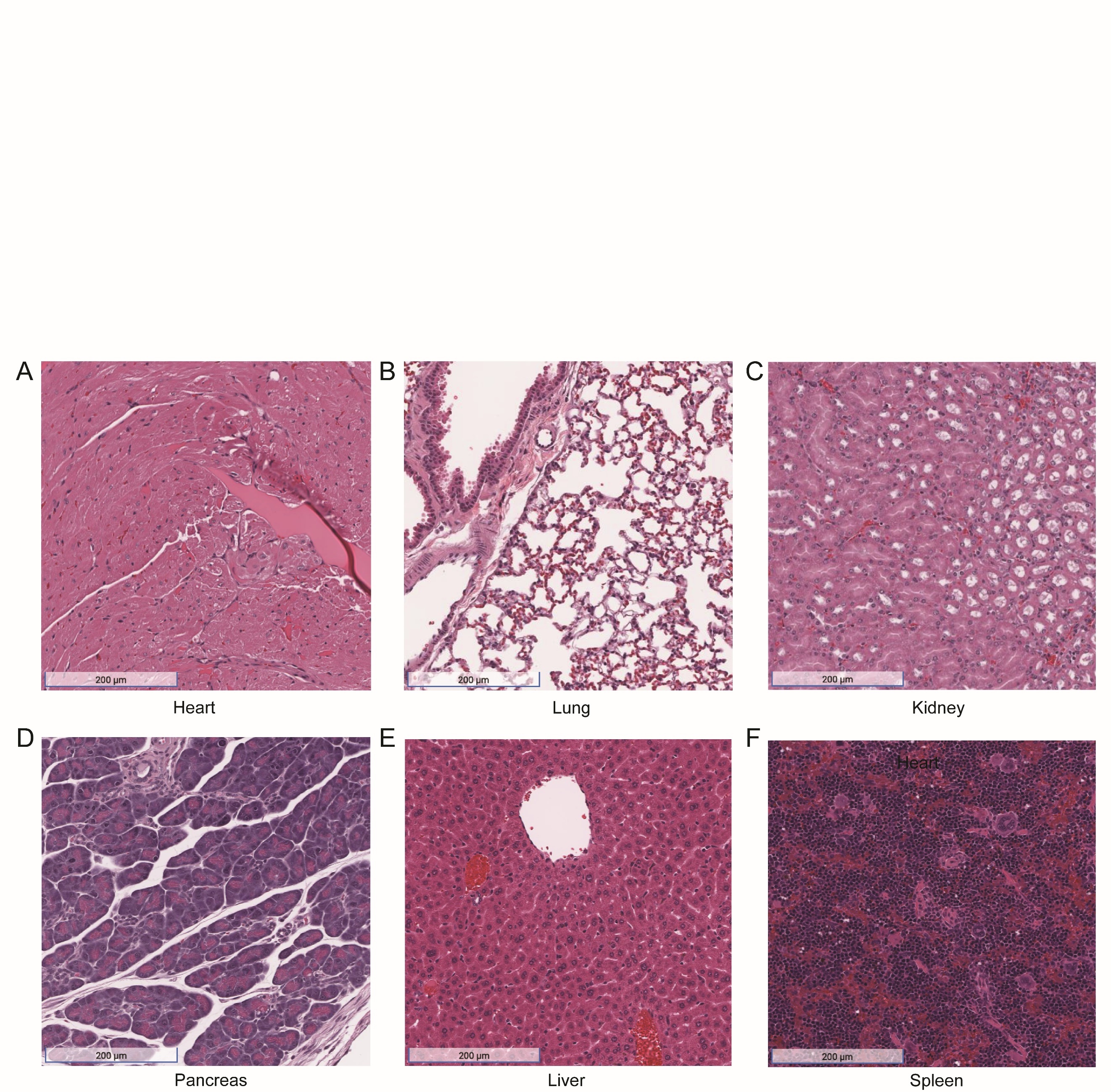

**Supplementary Figure 4.** Representative H&E images of organs of NSG mice treated with ME3BP-7 for 4 weeks: (**A**) heart, (**B**) lung, (**C**) kidney, (**D**) pancreas, (**E**) liver, (**F**) spleen x10

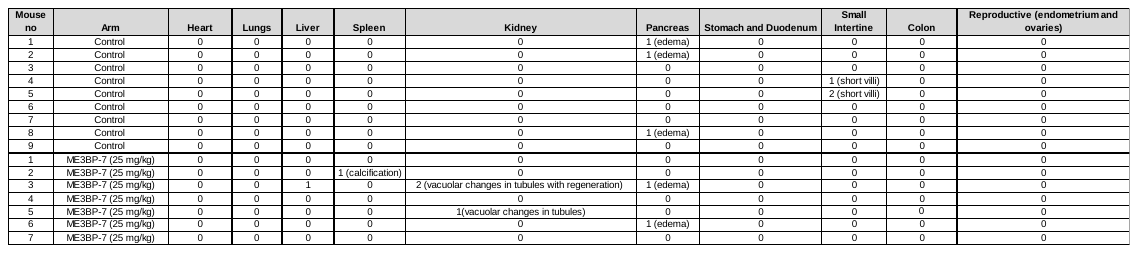

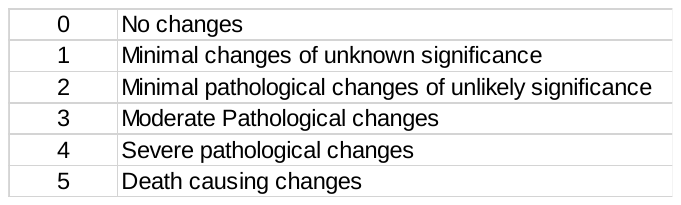

**Supplementary Table 1.** Gross pathology and assessment of potential tissue damage in organs of mice treated for 4 weeks (12 doses) with ME3BP-7

**Supplementary Figure 5.** ME3BP-7 reduces tumor burden in orthotopically implanted human patient derived xenografts from a PDAC metastatic site with focally expressed MCT-1 (TM01098). (**A**) Timeline and design of in vivo therapeutic study. (**B**) IHC sections of orthotopic PDx (TM01098) showing focal expression of (**C**) Representative Ultrasound image of orthotopically implanted tumors. (**D**) Mean tumor volumes of 7 and 10 NSG mice in each group as determined by ultrasound measurements on indicated days. (**E**) Weights of residual tumors harvested at end of therapy. (****** *P* < 0.01 Mann-Whitney U test)

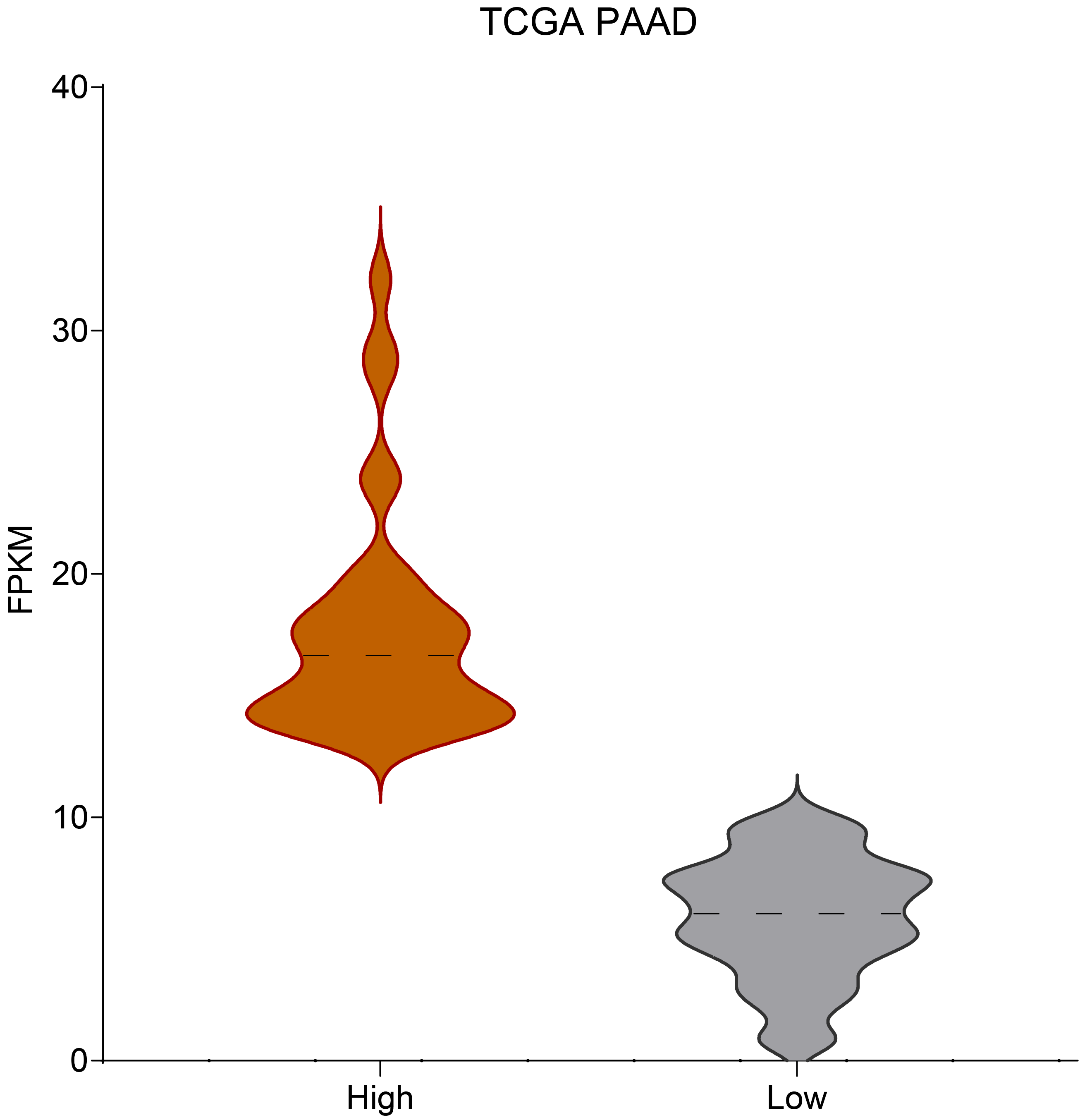

**Supplementary Fig. 6**: Violin plot of TCGA data for PDACs

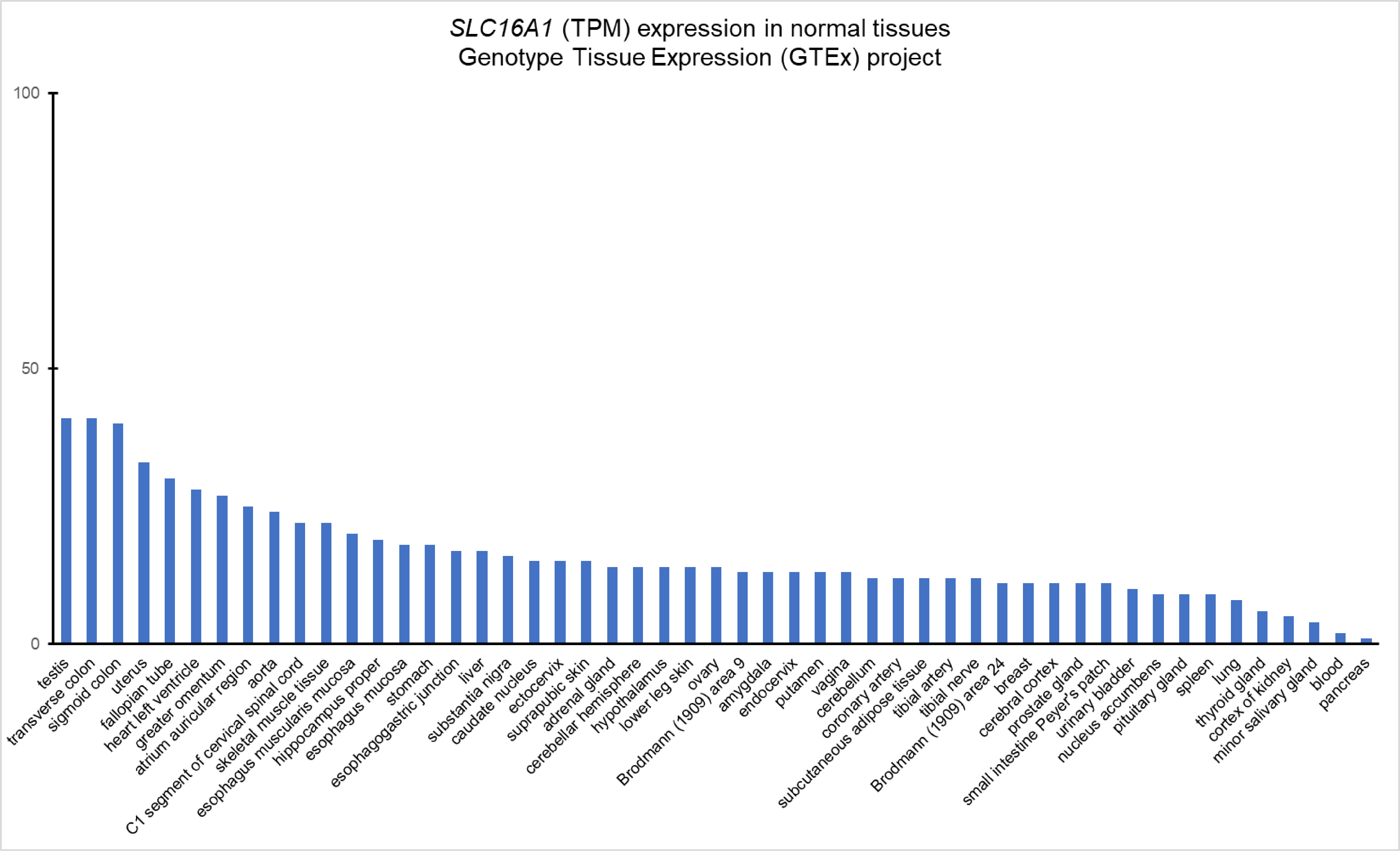

**Supplementary Fig. 7:** GTEx dataset showing expression of MCT-1 across 51 tissue types.

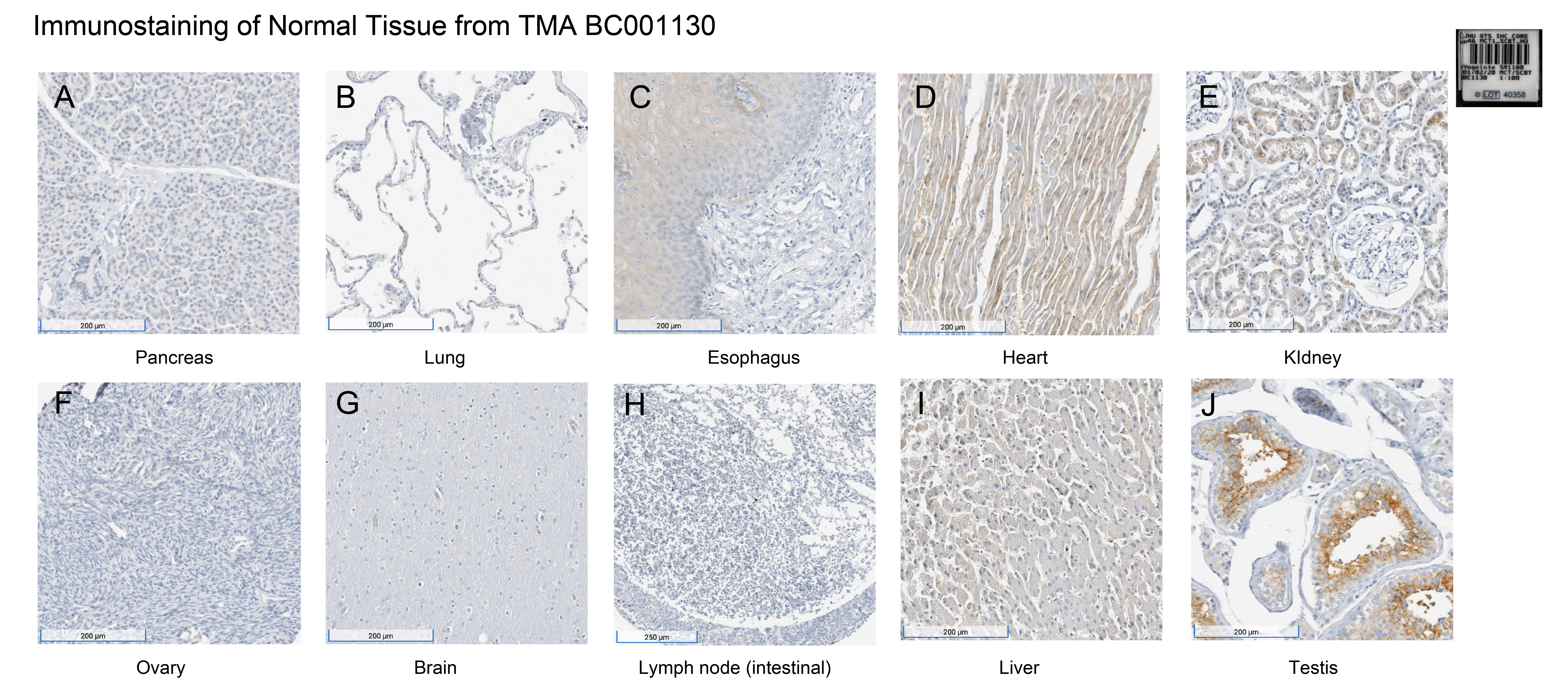

**Supplementary Figure 8**: IHC of normal tissues from human TMA BC001130 (10x)

(A) pancreas, (B) lung, (C) esophagus, (D) heart, (E) kidney (F) ovary, (G) brain, (H) intestinal lymph node, (I) liver and (J) testis
